# Trophic ecology of the African riverine elephant fishes (Mormyridae)

**DOI:** 10.1101/2023.06.07.543841

**Authors:** Gina Maria Sommer, Samuel Didier Njom, Adrian Indermaur, Arnold Roger Bitja Nyom, Petra Horká, Jaroslav Kukla, Zuzana Musilova

## Abstract

Multiple species of the elephant fishes (Mormyridae) commonly coexist in sympatry in most African tropical rivers and lakes. In this study, we investigated the trophic ecology and potential trophic niche partitioning of eleven mormyrid fish species from the Sanaga River system (Cameroon) using the stable isotopes of carbon and nitrogen of muscles and of trophic prey samples. Albeit mormyrids mainly feed on invertebrates, we found differences in isotope signals and the trophic niche partitioning in the studied species. We further show that species with elongated snout tend to show higher carbon and nitrogen isotope signals, suggesting a potential role of snout shape in their trophic preferences. Furthermore, we found significant differences in isotopic signatures within the *Mormyrus* genus, highlighting ecological niche diversification among three closely related species. We also report on different isotopic signals between seasons of the year in four species, possibly caused by species migration and/or anthropogenic agricultural activities. Overall, our research presents robust evidence of the trophic niche partitioning within the entire mormyrid species community, shedding light on the enigmatic evolutionary history of these fascinating African fishes.

## 1. Introduction

Elephant fishes, Mormyridae, exclusively occur in freshwater habitats of Africa, and they are well known for their additional electric sensory system. The produced electric pulses serve for electrolocation in turbid waters, as well as for conspecific communication (Sullivan et al., 2002; Carlson et al., 2011) and evolution of species-specific signals is often associated with mormyrid diversification (currently 228 described species; Fricke et al., 2023). Apart from their sensory systems, little is known about the general life history of mormyrids. Seasonal migrations have been reported in some species in several river systems (OKEDI, 1965; BLAKE, 1977, VAN DER WAAL, 1996), and seasonal changes are possibly impacting food availability and in some cases even shifts of diets and consequently trophic position of aquatic animals (e.g., NASIMA et al., 2020; MCMEANS et al., 2019). Studies on ecological aspects like movement patterns, reproductive behavior and feeding strategies are scarce in mormyrids revealing mostly complex behavioral traits such as pack hunting in *Mormyrops anguilloides* in Lake Malawi (Arnegard & Carlson, 2005) or the sound production of *Pollimyrus isidori* mediating their courtship (Crawford, Jacob & BenÉch, 1997). Feulner et al. (2007) investigated diversification of the genus *Campylomormyrus* and found that shape of the snout protrusions may be associated with different trophic specializations, although this needs further confirmation. Studies on trophic ecology show that mormyrids are mainly invertivorous (Winemiller & Adite, 1997; adjibade et al., 2019; Peel et al., 2019) or rarely piscivorous (*Mormyrops anguilloides*; Bailey, 1994; Skelton, 2001). Differences in trophic preferences have previously been described in co-existent species from lake Ayame, Ivory coast (KouamÉlan et al. (2006) or Sokoto-Rima river basin, Nigeria (Hyslop, 1986) based on the stomach content analyses showing that different species may specialize on different groups of insect larvae.

The method of stable isotope analysis is a powerful tool to illuminate trophic preferences. Heavier stable isotopes of carbon (^13^C), nitrogen (^15^N) or sulfur (commonly ^34^S) can be detected by stable isotope mass spectrometry and accumulate differently in the organisms compared to the common isotope: the lighter isotopes form weaker bonds than the heavier ones as well as tend to react faster (Jardine et al., 2003). Reports based on stable isotope analyses in mormyrids are scarce (Soto et al., 2019) finding just subtle differences among species compared to the effect of sexual selection (Arnegard et al., 2010) while being more commonly used in other African fishes (e.g. cichlids, Muschick et al., 2012).

In this study, we focus on the stable isotope analysis in eleven species of mormyrid fishes from the Sanaga River (Cameroon) to test for differences in trophic resources. We aim to provide the isotopic analysis throughout the entire mormyrid species community and shed light on the Cameroonian freshwater fish diversity in one of the less regulated rivers in Africa (until recently). We further focus on species collected at different seasons of the year (i.e., the wet and the dry season and a transition phase) and include fish samples from multiple localities on the main Sanaga River around the Nachtigal falls, as well as in some nearby tributaries.

## 2. Material and Methods

### 2.1. Sampling

The study site for this work was the Sanaga River in Cameroon, the country’s largest river of over 918 kilometers long (DUBREUIL et al., 1975). There are three large contiguous segments, Upper Sanaga, Middle Sanaga and Lower Sanaga (Bitja Nyom et al., 2020) and this study took place in the middle part of the Sanaga River. Sampling localities were located along the main stream in the proximity of the cascades of Nachtigal complemented by few inflow stream locations (tributaries) (see Map in Fig. 1). There exist different seasons in this Guinean equatorial climate zone (OLIVRY, 1986; AMOUGOU et al., 2015), namely a main wet season (from August until November) and a main dry season (from December to March) plus a transition between the dry and the wet season (from April until July), sometimes divided as a shorter wet season lasting from April until June and a shorter dry season (from July until early August) (BITJA NYOM & PARISELLE, 2015).The fish were caught using gillnets of different mesh sizes. After euthanizing the fish with a lethal amount of MS-222, photographs and measurements were taken, and fin clips collected for DNA analyses. Afterwards, a 1 to 2 cm piece of lateral muscle was dissected from an area between the ventral and the caudal fin in proximity to the lateral line. Eleven species (*Campylomormyrus phantasticus* (Pellegrin, 1927), *Hippopotamyrus castor* Pappenheim, 1906, *Marcusenius mento* (Boulenger, 1890), *Marcusenius sanagaensis* Boden, Teugels & Hopkins, 1997, *Mormyrops anguilloides* (Linnaeus, 1758), *Mormyrops caballus* Pellegrin, 1927, *Mormyrus tapirus* Pappenheim, 1905, *Mormyrus* sp. “short snout”, *Mormyrus* sp. “long snout”, *Paramormyrops batesii* (Boulenger, 1906) and *Petrocephalus christyi* Boulenger, 1920) were sampled in 2017 and 2018. The number of individuals per species vary and the representation of each season is not always complete and at times unequally distributed (Table 1). All fishes were collected with appropriate research permits namely N° 007, 048, 049/MINRESI/B00/C00/C10/C12 and N° 2376/PRBS/MINFOF/SETAT/SG/DAFP/SDVEF/SC/ENJ granted by the Ministry of Scientific Research and Innovation and the Ministry of Forestry and Wildlife in Cameroon.

**Figure 1:**
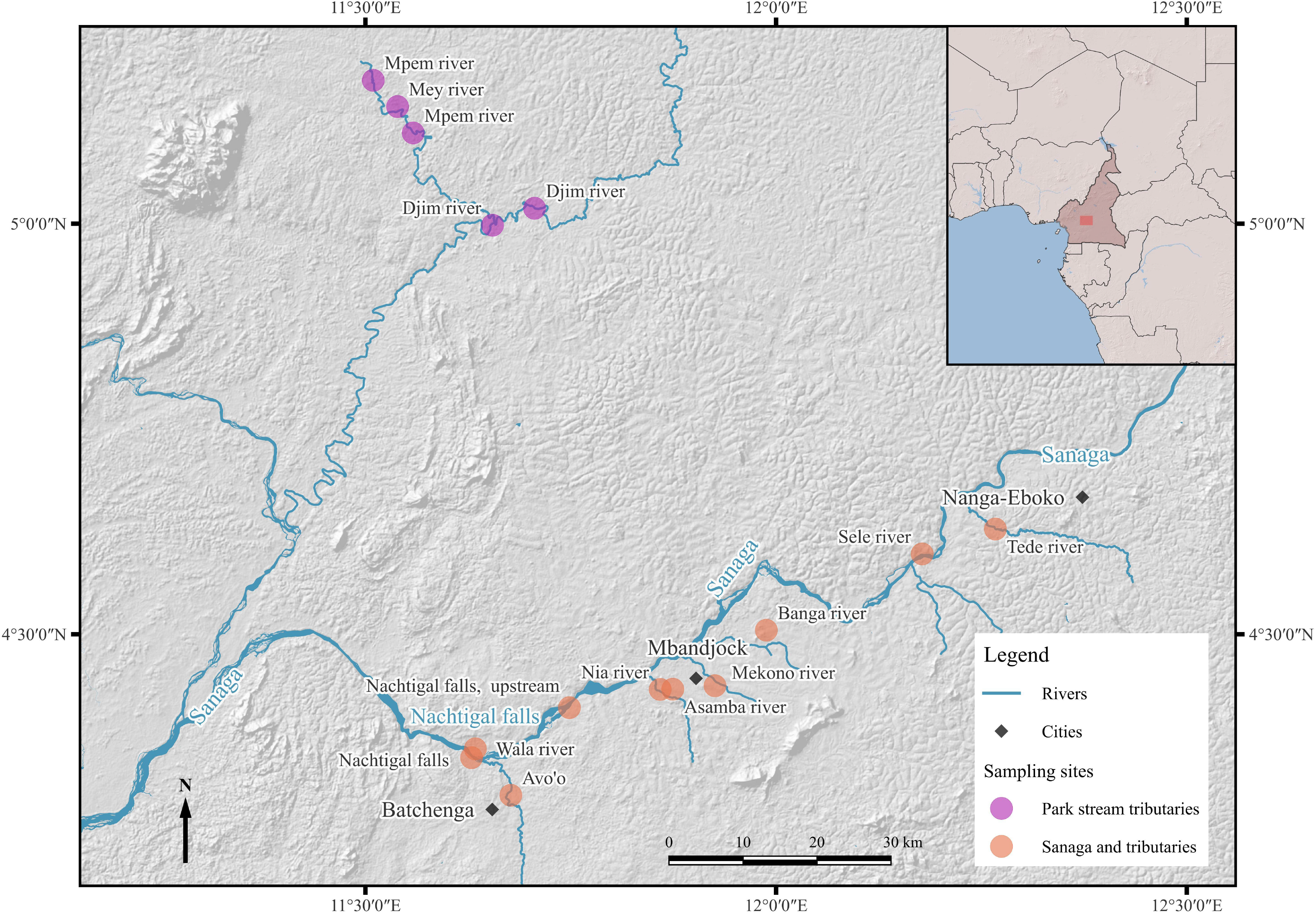
Overview of the study site: Map of the section of the middle Sanaga river close to the Nachtigal Falls (Chutes de Nachtigal) in Cameroon, Africa. The most frequented sampled sites are up- and downstream of Nachtigal falls and are in the vicinity of the town Batchenga (black dot). (Basemap: https://www.esri.com/en-us/maps-we-love/overview, borders: https://www.diva-gis.org/gdata, rivers, streams and places: openstreetmap.org).

**Table 1:**
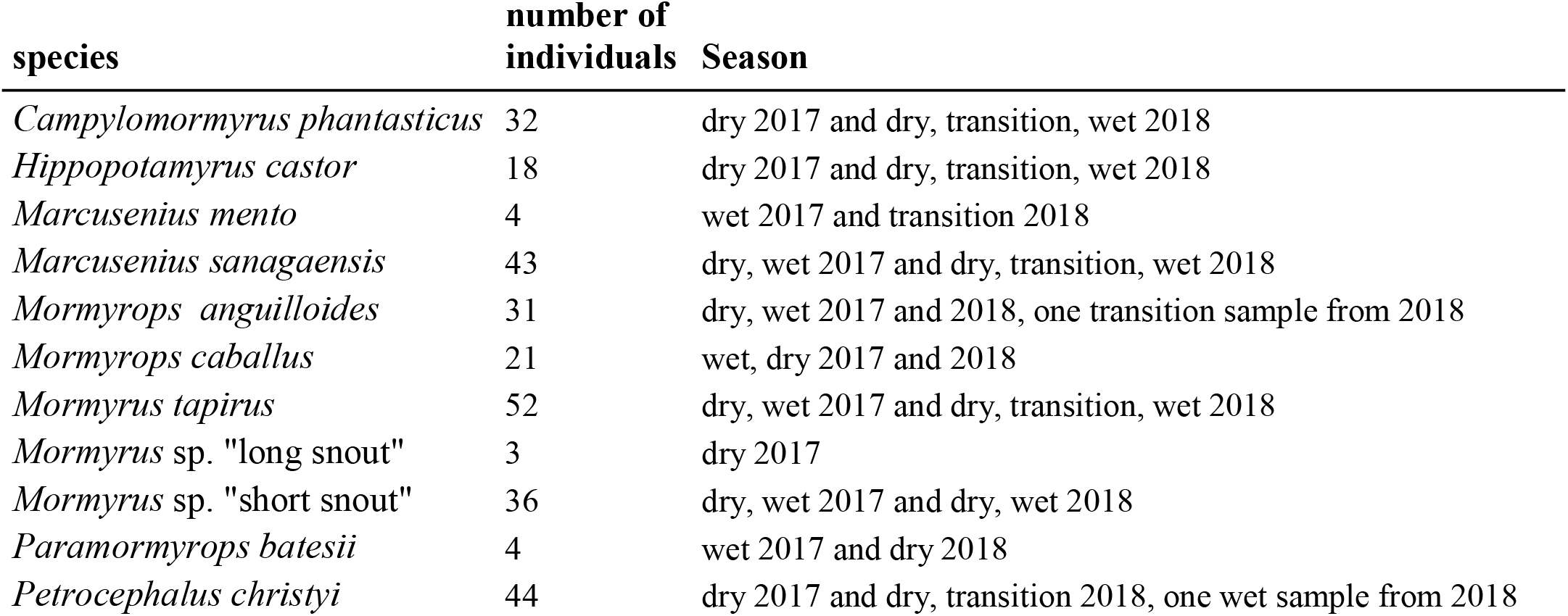
overview of collected species and their representation. Please note that *Mormyrus* sp. “short snout” and *Mormyrus* sp. “long snout” refer to so far undescribed species and thus have these preliminary names in this study.

In total we collected samples from 288 individuals of 11 species. All samples were transfered in 96% ethanol and stored in a freezer at −20 degrees. We further complemented our muscle sampling by adding trophic samples consisting of the individual items from the stomachs of nine individuals (two *M. caballus*, one *M. anguilloides*, two *H. castor*, two *M. tapirus* and one *M.* sp. “short snout”), as well as of benthic samples, i.e. plant material and invertebrates collected from the river bed area, the riverbank and the water column of the Sanaga River and a tributary, namely from Nachtigal falls and the Wala river. These samples target potential prey and a serve as a reference isotope background. The organisms from these environmental samples were identified taxonomically under a binocular microscope to the highest level possible, from sometimes only class (Gastropoda), mostly the order (e.g. Ephemeroptera) and seldom to a family level (e.g. Hydrophilidae, Elmidae) with the help of pictures taken from stomach content material and invertebrate identification guides (Moisan J. *et al*., 2010; Mary, N., 2017). The water plant material was not further identified, and the various plants were mixed to achieve an amount of over 2 mg and labelled accordingly as the group called “plant”. The isotope values generated by both, the stomach content material and the benthic material, are jointly represented when later referring to the values of the “trophic samples”, unless otherwise specified.

### 2.2. Sample processing

The muscle tissue samples were transferred to 2ml Eppendorf tubes and were dried for 48 hours at 60 °C or at 45 °C for 72 hours over the weekend in a drying oven. If bone structures were present in the sample, they were carefully removed with a clean forceps. Then samples were ground with two beads at the frequency 25/s for 20 seconds or at 30/s for 30 seconds in case of a bigger piece using a ball mill (MM400, Retsch, Germany). After taking the beads out, one millilitre of ethanol was added, and sample has been shaken until homogenous. The samples were then centrifuged for 30 seconds at 6000 rpm. About 600 microliters of ethanol were taken out with a pipette and a second drying step was followed at 24 up to 48 hours for 60 °C. In case the sample was not completely transformed into the powder yet, an additional ball grounding was performed at 25/s frequency for 10 seconds and then centrifuged for 30 seconds at 6000 rpm and final drying for 15 minutes and 60 °C. Subsequently, the samples were weighted in tin capsules to achieve 0.5 milligrams ideally while allowing a range of 0.45 to 0.55 mg and folded into tiny cubes using a thin and thick tweezer and put in labelled sample boxes. The trophic samples were processed in the same manner except grounding happened partly manually and sometimes with only one silver ball. Plant material had to achieve a weight of 3 mg. Some trophic samples were too light and had to be excluded from further analysis (17 samples out of 64). Total carbon and nitrogen amounts as well as the isotopic ratios δ^13^C and δ^15^N were measured with a Delta V Advantage mass spectrometer coupled to Conflo IV and elemental analyzer Flash 2000 using a Thermal Conductivity detector (all instrumentation by Thermo Fisher Scientific, Bremen, Germany). The carbon and nitrogen isotope ratios are expressed following the delta notation, which is the following: δX = 1000*(Rsample/Rstandard – 1), where X stands for ^13^C or ^15^N, respectively, and R is the relative amount of carbon or nitrogen isotopes (R□=□^13^C/^12^C or ^15^N/^14^N). International standards were applied to normalise the isotope ratios: The Vienna Pee Dee Belemnite (VPDB) scale (Coplen, 1996) and atmospheric N2 was used for the carbon and the nitrogen isotope ratios, respectively. In addition to repeated measurements of a series of international standards (IAEA-CH3, IAEA-CH6, IAEA-600, IAEA-N1, IAEA-N2, IAEA-NO3), a glycine standard was included after every 10th sample to provide calibration for elemental composition as well as serve as a quality control for isotopic measurement. Analytical precision resulted within ± 0.2 ‰ for both δ^13^C and δ^15^N.

### 2.3. Data analysis

Delta 13C (δ^13^C) and delta 15N (δ^15^N) values were analysed in R Studio Statistical Software (version 1.4.1717; RStudio Team (2021). RStudio: Integrated Development Environment for R. RStudio, PBC, Boston, MA URL http://www.rstudio.com/.). The package “tRophicPosition” (Quezada-Romegialli C. *et al*., 2018) was used furthermore to access the trophic position of the investigated species using one baseline of all benthic sourced samples from the trophic samples. The used trophic discrimination factors (TDF) were McCutchan (McCutchan’s *et al*. 2003) for a fish muscle tissue which means a mean of 2.9 ‰ ± 0.32 se for δ^15^N and 1.3 ‰ ± 0.3 se of for δ^13^C. Tropic position (TP) was estimated using a λ (lambda) = 2 for using also secondary consumers as baseline as no primary producer was available. The conventional range of trophic levels ranges from 1 (primary producers) to 5 (2 to 4.5 for fishes in freshwater ecosystems) (Trites, 2001; YoĞurtÇuoĞlu et al., 2020). The plant material collected were several species but had to be pooled together to increase weight and thus has only informative value. The model was run with 4 chains and for 20000 iterations. To access the significance of the differences observed in muscle isotopic signals mean values between different seasons one-way ANOVAs as well as Tukey tests to judge the results of the ANOVA were conducted. Alpha was set to 0.01 and the critical value q was found accordingly in available tables online (see Table 3).

## 3. Results

### 3.1. Muscle samples analysis

In total, we analysed 288 samples of 11 species of the mormyrid community in the Sanaga River, and we found that the stable isotope signals partly differ among the species (Fig. 2). The *Marcusenius* species and *Paramormyrops batesii* share lower δ^13^C and δ^15^N values in comparison to all other species. Interestingly, some species from different genera possess similar isotopic signatures like *Hippopotamyrus castor* and *Mormyrus tapirus,* or *Mormyrops caballus* and *Campylomormyrus phantasticus* whereas the species from the same genus are more separated from each other. This is especially obvious in the *Mormyrus* genus (Figs 2-3).

**Figure 2:**
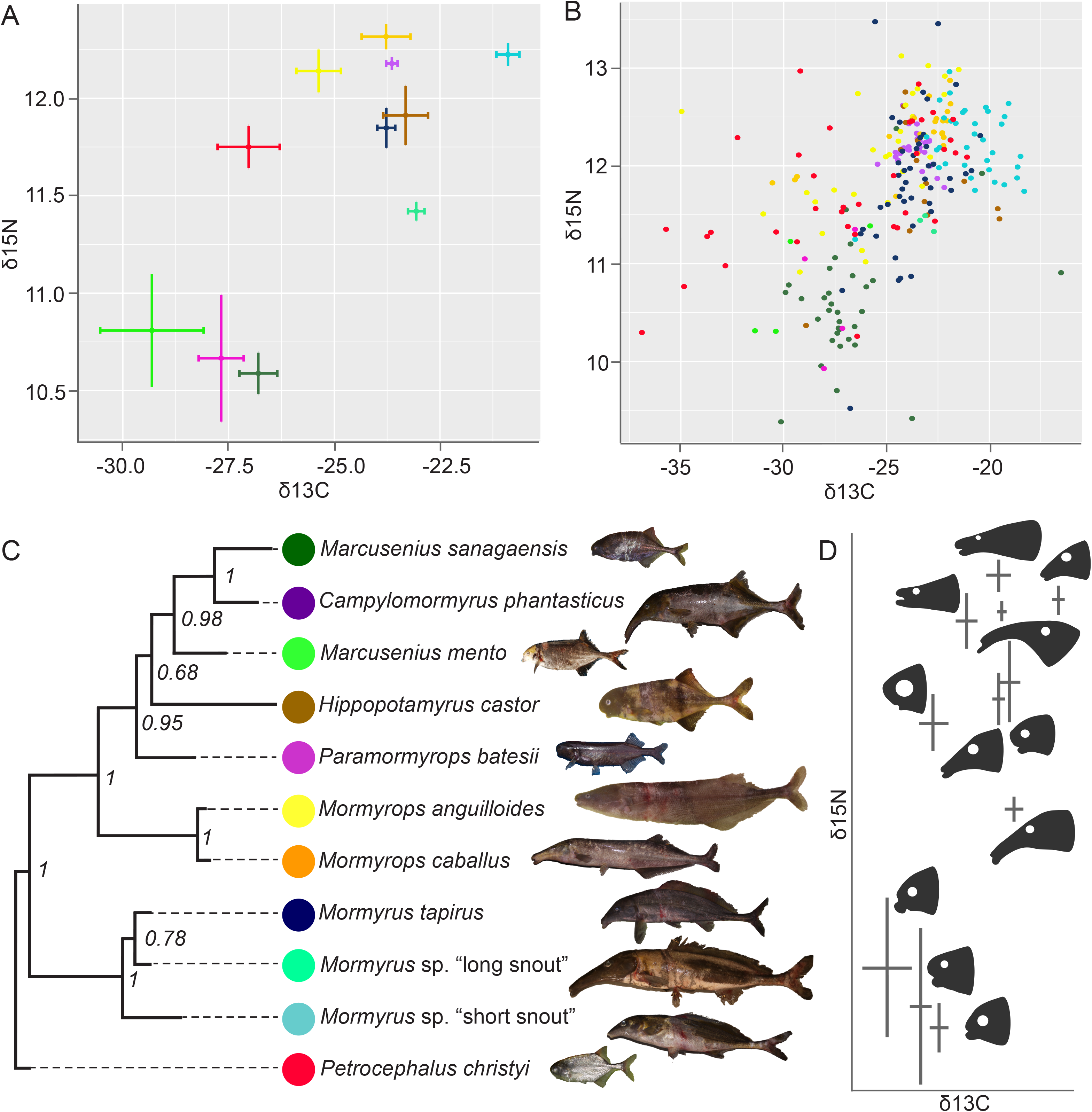
Stable isotope signature of eleven Sanaga river mormyrid species. A) Results of the stable isotope analysis of the muscle samples - mean and standard errors shown. δ^15^ nitrogen values on the y-axis and δ^13^ carbon values on the x-axis; unit in parts per thousand (‰). B) Stable isotopic signatures shown for all individuals. Note that the plots do not include the transition season samples specifically addressed in the text and in Figure 5. Note the different scale of plot A and B. C) Phylogenetic relationships of the studied mormyrid species based on the COI sequences and reconstructed by Bayesian inference. The colour code corresponds to the species in the plots above. D) Head shape of different species projected on the results of the stable isotope analysis (plot A) to visualize that species with snout protrusion tend to exhibit higher values in δ^13^ C and/or δ^15^ N.

**Figure 3:**
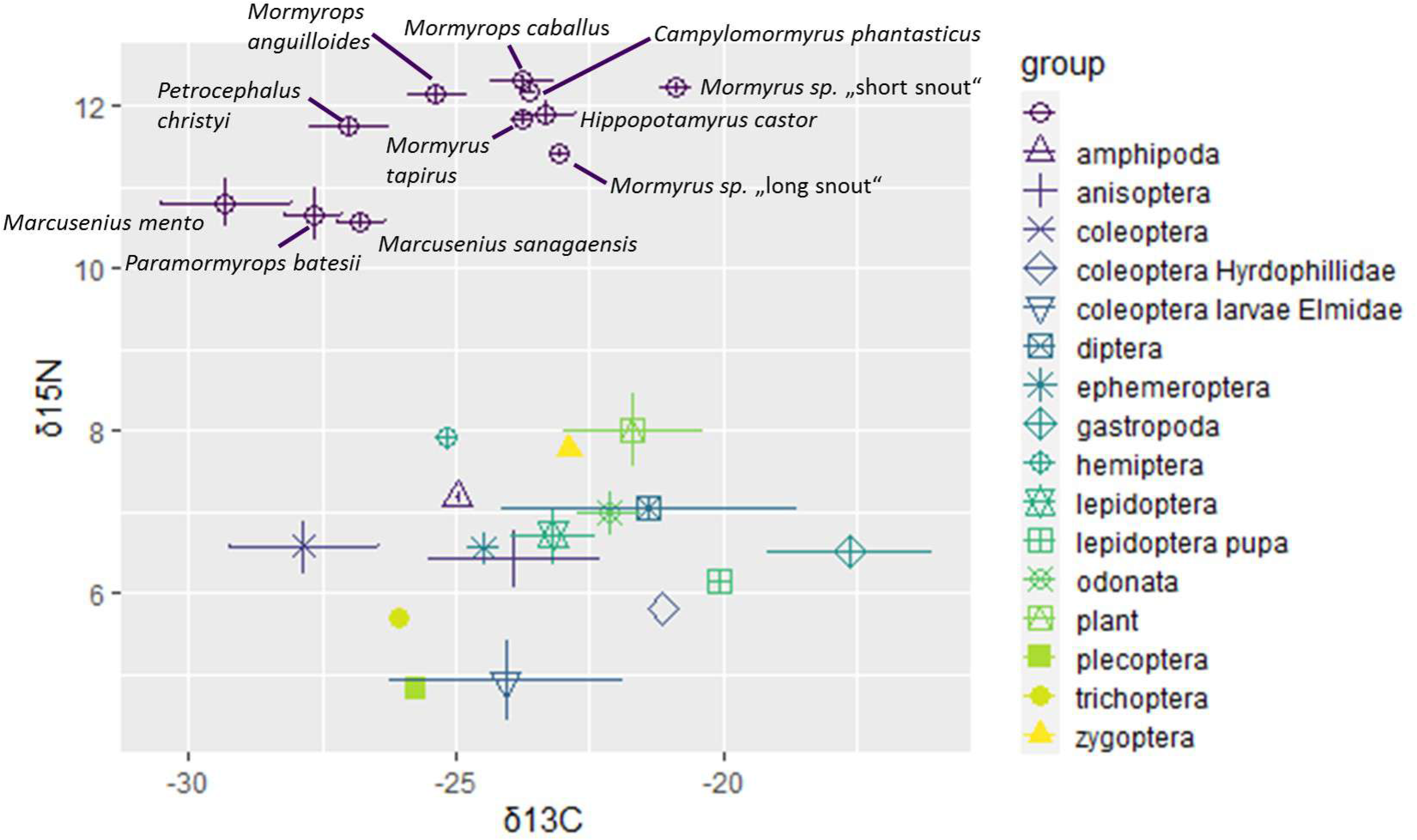
combined isotopes plot showing both the muscle isotope means and standard errors together with the trophic samples values. δ^15^ nitrogen values on the y-axis and δ^13^ carbon values on the x-axis; unit in parts per thousand (‰).

### 3.2. Trophic samples

As a baseline, we analyzed trophic samples of possible prey items, i.e. selected items from the fish stomachs as well as benthic samples. We analysed the stable isotope values of 16 potential prey groups. Some individual samples had to be pooled together in order to reach the threshold weight needed for the analysis, like the group Plecoptera, Trichoptera and Ephemeroptera. The δ^15^N values range from 4.9 to 5.0 parts per thousand (‰) and the δ^13^C values from −17,5 to −28 parts per thousand. Gastropoda has the highest δ^13^C values and Coleoptera the lowest. The plant material, Zygoptera and Hemiptera display the highest δ^15^N values whereas Plecoptera and Elmidae (Coleoptera) the lowest. Only three groups (Anisoptera, Gastropoda and Elmidae (Coleoptera)) were simultaneously present in the stomach samples and the benthic items. The isotopic signatures did only marginally differ, mainly in Elmidae (Figure S1 for the benthic items and Figure S2 for the stomach samples). We noticed expected pattern of higher trophic position (δ^15^N) in mormyrid fish muscles compared to the prey items (Fig. 3).

### 3.3. Calculations of trophic position

The analysis performed with the tRophicPosition R package (QUEZADA-ROMEGIALLI C. et al., 2018) with a benthic prey baseline revealed the trophic positions of all mormyrid species between 3 and 4. Pairwise comparisons within genera revealed only marginal differences in the trophic position of two species of *Mormyrops* (mean TP of 3.8 vs. 3.7) (Fig. 4A) and *Marcusenius* (3.612 vs. 3.545) (Fig. 4B). On the other hand, we found differences among the three *Mormyrus* species. Namely, *Mormyrus* sp. “short snout” shows a notable difference (3.912) from *Mormyrus* sp. “long snout” (3.605) and from *Mormyrus tapirus* (3.772; Fig. 4C,D,E). Accordingly, statistical analysis (t-tests) revealed significant differences of the stable isotope values only in the *Mormyrus* species (see Table 2 below). This is in accordance with the nearly congruent trophic positions of the *Mormyrops* and the *Marcusenius* species but also reflects the separation of *Mormyrus* sp. “short snout” from *Mormyrus tapirus* and from *Mormyrus* sp. “long snout” nicely.

**Figure 4:**
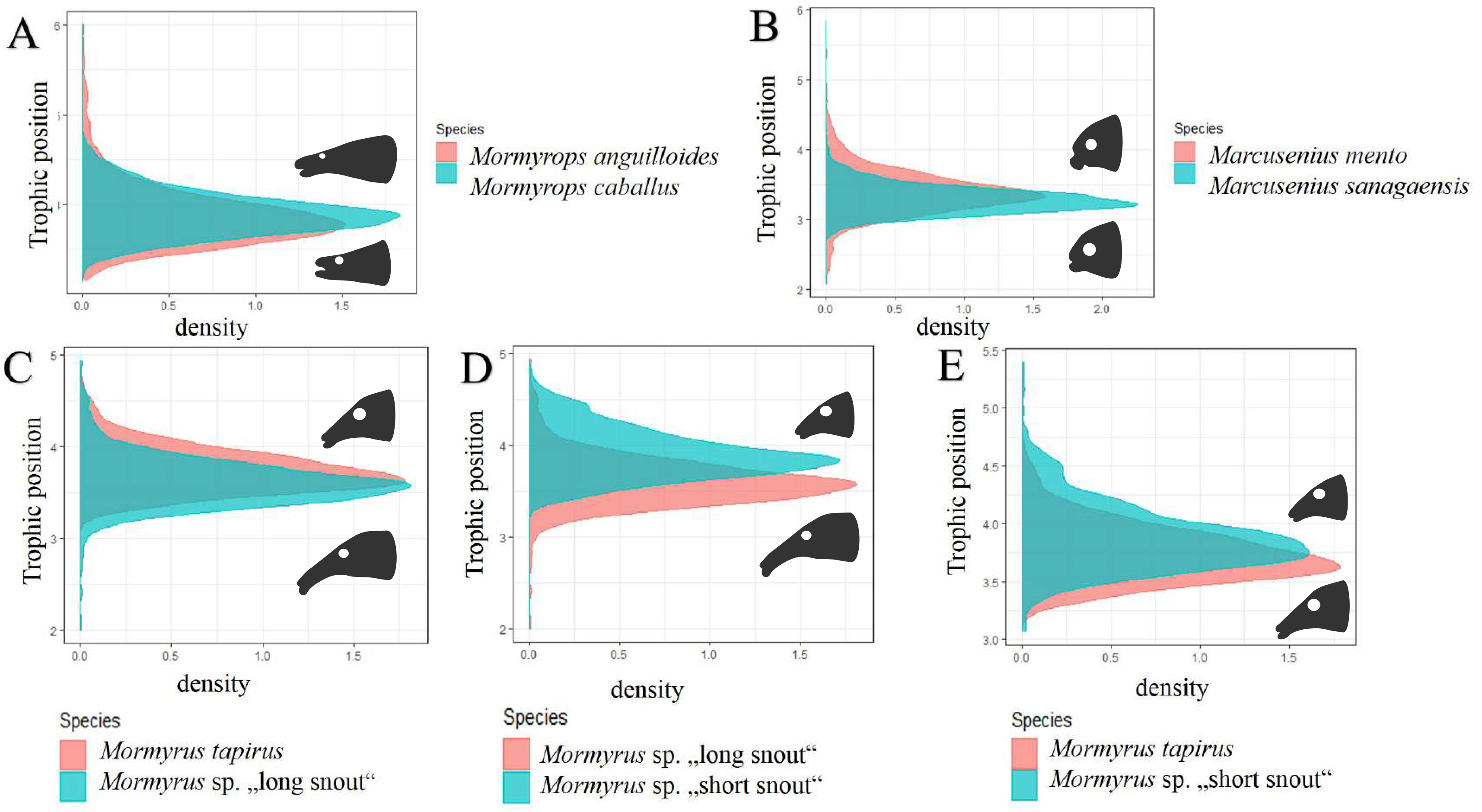
TP (Trophic Position) plots comparing TPs between two species within one genus. A) *Mormyrops anguilloides* vs. *Mormyrops caballus*, B) *Marcusenius sanagaensis* vs. *Marcusenius mento*. C)-E) Three species of *Mormyrus* show larger differences among themselves, and in this case species with shorter protrusion tend to have a higher trophic position. C) *Mormyrus tapirus* vs. *Mormyrus* sp. “short snout”, D) *Mormyrus* sp. “long snout” vs. *Mormyrus* sp. “short snout”, E) *Mormyrus tapirus* vs. *Mormyrus* sp. “long snout”. The baseline was generated from all benthic-sourced samples from the trophic samples.

**Table 2:** t-tests performed for delta nitrogen values as well as delta carbon values for species comparisons of the same genus.

| **t-test with different variances** | **p-value**<br>**(two-tailed)** |
| --- | --- |
| $\delta^{13}\text{C}$ *Mormyrops caballus* vs. *Mormyrops anguilloides* | 0.116 |
| $\delta^{15}\text{N}$ *Mormyrops caballus* vs. *Mormyrops anguilloides* | 0.650 |
| $\delta^{13}\text{C}$ *Marcusenius mento* vs. *Marcusenius sanagaensis* | 0.126 |
| $\delta^{15}\text{N}$ *Marcusenius mento* vs. *Marcusenius sanagaensis* | 0.515 |
| $\delta^{13}\text{C}$ *Mormyrus tapirus* vs. *Mormyrus* sp."short snout" | 4.42E-12* |
| $\delta^{15}\text{N}$ *Mormyrus tapirus* vs. *Mormyrus* sp."short snout" | 0.002* |
| $\delta^{13}\text{C}$ *Mormyrus tapirus* vs. *Mormyrus* sp."long snout" | 0.04 |
| $\delta^{15}\text{N}$ *Mormyrus tapirus* vs. *Mormyrus* sp."long snout" | 0.002* |
| $\delta^{13}\text{C}$ *Mormyrus long "snout"* vs. *Mormyrus* sp."short snout" | 1.471E-05* |
| $\delta^{15}\text{N}$ *Mormyrus long "snout"* vs. *Mormyrus* sp."short snout" | 1.003E-05* |

\* indicates a significant p-value (p-value < 0.025)

### 3.4. The impact of the season

We analyzed samples from different seasons, namely during the dry season (February and March), the wet season (August, September, October and November) and the transition phase between these two seasons (in the months of May, June and July). We investigated the influence of the season on the stable isotope values and found no difference when comparing dry and wet season values. We did find differences though between the samples collected during the transition phase and the two main seasons. Albeit the low number of samples from the transition phase (only for five species; Fig. 5), four species show significant differences between the wet vs. transition and/or dry vs. transition phase (Table 2). The one-way ANOVAs and the subsequent Tukey-Kramer tests revealed that in two species, *Mormyrus tapirus* and *Marcusenius sanagaensis*, both stable isotopes δ^15^N and δ^13^C were significantly different between the seasons (pairwise comparisons of wet-vs.-transition and dry-vs.-transition). In *Campylomormyrus phantasticus* a difference was found only in the δ^13^C signal for the dry-vs.-transition samples, while in *Petrocephalus christyi* the δ^15^N values are different, although the statistical analysis could not be performed for all pairs (only one wet-season sample was available). Only in *Hippopotamyrus castor* no differences in the stable isotope signal was noticed. (Table 3).

**Figure 5:**
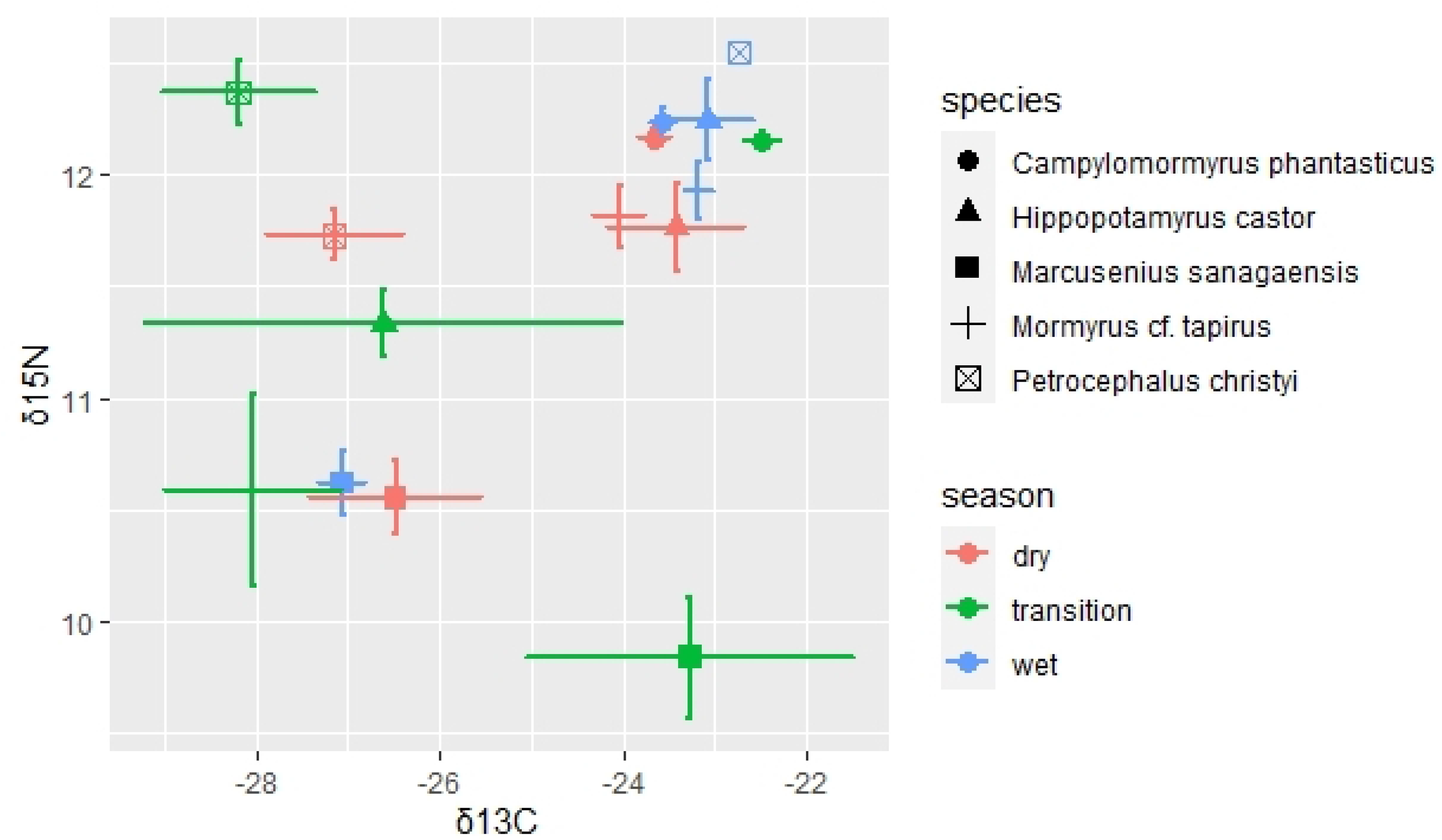
season impact plot of stable isotopes: plotted means and standard errors of five mormyrid species separated by wet (blue-coloured), dry (red-coloured) and transition (green-coloured) season. δ^15^ nitrogen values on the y-axis and δ^13^ carbon values on the x-axis; unit in parts per thousand (‰). Note that the *Petrocephalus christyi* wet sample is only one sample and thus no standard errors are available. The dry season is encompassing the months of February and March, the wet season the months August, September, October and November and the transition season the months of May, June and July.

**Table 3:** results of single-factor ANOVAs and Tukey-Kramer test (if applicable) for all species with transition season samples.

| single-factor ANOVA | p-value | df | Tukey-Kramer q-stat values | dry vs wet | dry vs transition | wet vs transition | critical value with $\alpha=0.01$ |
| --- | --- | --- | --- | --- | --- | --- | --- |
| *Mormyrus tapirus* $\delta^{13}\text{C}$ | 1.288E-07* | 50 | | 2.476 | 8.353** | 9.242** | 2.678 |
| *Mormyrus tapirus* $\delta^{15}\text{N}$ | 0.001* | 50 | | 0.677 | 5.253** | 5.233** | 2.678 |
| *Marcusenius sanagaensis* $\delta^{13}\text{C}$ | 0.027* | 40 | | 0.669 | 3.157** | 3.861** | 2.704 |
| *Marcusenius sanagaensis* $\delta^{15}\text{N}$ | 0.011* | 40 | | 0.382 | 3.796** | 4.268** | 2.704 |
| *Petrocephalus christyi* $\delta^{13}\text{C}$ dry vs transition | 0.520 | 19 | | | | | |
| *Petrocephalus christyi* $\delta^{15}\text{N}$ dry vs transition | 0.011* | 19 | | | | | |
| *Campylomormyrus phantasticus* $\delta^{13}\text{C}$ | 0.016* | 31 | | 0.378 | 4.354** | 3.53** | 2.750 |
| *Campylomormyrus phantasticus* $\delta^{15}\text{N}$ | 0.654 | 31 | | | | | |
| *Hippopotamyrus castor* $\delta^{13}\text{C}$ | 0.232 | 14 | | | | | |
| *Hippopotamyrus castor* $\delta^{15}\text{N}$ | 0.169 | 14 | | | | | |

\*indicates a p-value below 0.05 ; \*\* indicate q-stat value > critical value

## 4. Discussion and Conclusions

### Trophic niche partitioning in mormyrid fishes

We provide the first comprehensive overview of the mormyrid stable isotope signals at the community level. Our trophic ecology results are in accord with previous evidence of mormyrids feeding mostly on insect larvae (Van der Waal, 1985; Winemiller & Adite, 1997). There are, however, noticeable differences among species, such as the two *Marcusenius* species and *P. batesii* having a lower trophic position than the other mormyrids. We also observe that the mean isotopic signal of different species within one genus often differs more than when compared across other genera. This is the case for *Mormyrops caballus* and *Mormyrops anguilloides* with distinct signals in both δ^13^C and δ^15^N and more strikingly among the three species of the *Mormyrus* genus. *Mormyrus tapirus*, and two formally undescribed species, *M.* sp. “long snout”, and *M.* sp. “short snout” differ in δ^13^C and δ^15^N signals, and, accordingly, *Mormyrus* sp. “short snout” also differs from the other two other *Mormyrus* species in the trophic position analysis (Fig. 4). Our findings therefore suggest different trophic specializations for these closely related species. Trophic niche partitioning among closely related species from the same habitat has been commonly observed in African fishes (e.g. Nagelkerke, 1997; PolÁČik et al., 2014), while resource partitioning has been a general question in ecology since a long time (Hardin, 1960; Shoener, 1974). Stable isotope data have already proven to be helpful to visualize trophic niche partitioning within closely related fish species (Wang et. al, 2022) and our study is the first case of stable isotope analysis showing trophic niche partitioning in mormyrids. Differences in trophic preferences can be further observed on the jaw/mouth shape, which is often adapted to its specific purpose. For example, in croakers (Sciaenidae; Wang et al., 2022) the species with the most unique isotopic signatures also has a unique feeding strategy associated with different jaw shape (Zhang et al., 2020). Based on our results, mormyrid species from Sanaga with snout protrusion tend to show higher δ^13^C and/or δ^15^N signals when compared to species without elongated snouts, or, from another angle, species with higher δ^15^N values (∼ trophic level) show higher variability in the snout elongation and shapes (Fig. 2D). Notably, the three *Mormyrus* species have different isotopic signals and at the same time express anatomical differences in their trunk shape and jaw and lip morphology. *Mormyrus* sp. “short snout” has a more truncated upper lip protrusion, while *M. tapirus* has a longer lower lip, and *M.* sp. “long snout” has much longer snout than the two other *Mormyrus* species (Fig. 2). The overall mouth shape and orientation probably reflect subtle differences in microhabitat and prey preferences in these species. Our data therefore illustrates the differential trophic specialization associated with the snout shape differences even on the shorter evolutionary scale in the three closely related species. This seems to be in accord with the hypothesis of Feulner et al. (2007) who suggested possible variations in trophic ecology based on morphological differences in the snout protrusions in the *Campylomormyrus* genus from the lower Congo River (Feulner et al., 2007).

The trophic position analysis is mostly based on the δ^15^N values since it has been shown that trophic position is corresponding more to the trophic fractionation of δ^15^N than to δ^13^C (Post, 2002). The environmental benthic samples were set as a baseline for the analysis and these samples represent possible sources of carbon and nitrogen from the environment. We also tested the trophic position with the stomach samples with very similar results (see Supplementary material S3). Ideally, a primary producer is used as a baseline for such analysis, unfortunately it was not available in our study as the collected plant material was insufficient. Some authors have also claimed that macrophytes, the main primary producers in many aquatic ecosystems, were revealed to contribute less to aquatic food webs than assumed (Hamilton et al., 1992; Bunn & Boon, 1993; Forsberg et al., 1993; Thorp & Delong, 1994; Thorp et al., 1998; Lewis et al., 2001) or may have only local effects (Grosbois et al., 2017a) (Peel et al., 2019). Working on an understudied group of tropical fishes, such as mormyrids, may be challenging as other factors contribute to the final patterns. In our case, seasonal changes (possible migration), differing turnover rates, or unknown discrimination factors for Bayesian analysis. Turnover rates are generally tissue-dependent and are slower in muscle tissue or bone collagen (two to three months) than in plasma or blood (few days) (e.g., Madigan et al., 2021, Polischuk et al., 2001) but there are hints that the turnover rates can also differ among species, fish size and life stage (Bosley et al., 2002; Gaston et al., 2004; Madigan et al., 2012 and Madigan et al., 2021). For discrimination factors (i.e., the exact isotopic shift between prey and consumer), we used McCutchan’s (McCutchan et al. 2003) values for fish muscles with a mean of 2.9 ‰ ± 0.32 se for δ^15^N and 1.3 ‰ ± 0.3 se of for δ^13^C, although values of the trophic shifts differ especially for N depending on diet. Interestingly, McCutchan et al. (2003) found lower ^15^N trophic shift mean values (around 1.4‰) for consumers feeding exclusively on invertebrates in contrast to consumers of high-protein diets. In our study, we found the trophic discrepancy for δ^15^N at a minimum of 2.5‰ between mormyrid consumers and trophic samples (Fig. 3) and this could speak for a more high-protein diet in some mormyrid species. In order to reveal the trophic resources and more details on mormyrid prey, we suggest that our results could be complemented by a long-term stomach content analyses in the future as the two methods can complement each other (Davis et al., 2012; Layman et al., 2012; PolÁČik et al., 2014; Dillon et al., 2021).

### Effect of seasonality

Dry and wet seasons in tropical regions can be reflected in fish behaviour, migration, reproduction or food/prey availability. Impact of the season on the stable isotope values has been reported from multiple fish groups (Nashima et al., 2020; McMeans et al., 2019; Taylor, 2016 [doctoral thesis]; Wantzen et al., 2002) and seasonal variability seems to be especially relevant for partly flooded aquatic systems (Taylor et al., 2017; Merron & Mann, 1995; De Almeida et al. 1997; Wantzen et al. 2002; Eggleton & Schramm, 2004). The isotopic signal can be influenced by various factors, not only by the actual changes in the prey items. Some studies reported on fish to become more generalized feeders during the wet season (Costa-Pereira et al., 2017; Pool et al. 2017) and there are hints for increasing trophic position during the dry season (Wantzen et al. 2002), whereas other studies showed diverse foraging strategies with no universal seasonal influence pattern (Novakowski et al. 2008), (McMeans et al., 2019). In this study, we have specifically focused on the stable isotope signatures in different seasons throughout the year (Table 1 and S1). We found differences in isotopic signal between the transition phase (samples from May-July) compared to the wet season (August-November) or the dry season (February-March), whereas no difference was found between dry and wet season. The different isotope signal is present in four out of five tested species, i.e., in *Mormyrus tapirus*, *Marcusenius sanagaensis, Petrocephalus christyi* and *Campylomormyrus phantasticus,* while *Hippopotamyrus castor* did not show any differences throughout seasons. Therefore, the transition period seems to influence the diet of some, but not all mormyrid species. The habitats of the Sanaga River and tributaries are characterized by seasonal flooding during the wet season and are exposed to a rising water level already in the transition phase. Multiple factors could contribute to the seasonality differences. First, species seasonal migration could cause that individuals sampled in the transition phase may originate from different population than the dry- and wet-season samples. In general, mormyrids are known to migrate (Van der Waal, 1996; Olopade, 2013), mostly in relation with the wet seasons spawning (Okedi, 1965; Blake; 1977) but little is known for the tested species from the Sanaga River. At least one of the species, *Marcusenius sanagaensis,* is known to be more residing in the tributaries than in the main stream (personal observations on the study by Njom and others during several years of fieldwork). An alternative explanation would be in the turnover rates. If turnover rates are a few months for white muscle tissue like in most other more intensively studied fish species (Taylor et al., 2017), it could mean that a change of diet at the very beginning of the dry season and during the dry season would affect the transition phase values. Interestingly, this could be an anthropogenic induced cause due to the phytosanitary treatment of the plantations along the tributaries carried out towards the end of the major dry season (Njom et al., 2022). Such unusually large terrestrial input to the water may possibly impact the signal of some (but not all) species. Lastly, we can also explain the pattern partially by different sampling locations in the transition period for some species (Table S1). For example, the samples of *Marcusenius sanagaensis* from the Mekono River (a tributary with a pollution impact largely increased during the dry season) are different from the main Sanaga River and this seems to influence the results more than the season in this species (revealed by ANOVAs shown in Table S3). In *Mormyrus tapirus* both season and locality seem to both play a role. The difference in δ^13^C values (but not in δ^15^N) remains after exclusion of the most different samples are from Mpem River (as seen in the Supplementary table S3). Interestingly, the transition samples from the same locality (Mpem River) are not significantly different in *Marcusenius sanagaensis*, which illustrates again differences among species. Some differences of the transition samples, e.g. δ^13^C values of *Campylomormyrus phantasticus* (all samples are from one locality) or the δ^15^N values of *Petrocephalus christyi* remain irrespective of locality issue. Unfortunately, our sampling is not robust enough to test properly the effects of all aforementioned factors possibly involved in the seasonally different signal of the stable isotopes in mormyrids.

To conclude, here we present the first comprehensive study applying stable isotope analysis on mormyrid fishes at a species community level. We found differences in the isotopic signal among eleven species corresponding to putative feeding specialization and we also report on putative role of the snout shape in the trophic ecology of different species. We have specifically focused on the difference between three species of the *Mormyrus* genus with the clear trophic niche differentiation after the recent evolution of these species. We further report on seasonal differences in the isotopic signal in four species. Our study attempts to shed light on the biology of the evolutionarily successful and diverse - and yet understudied – group of tropical freshwater fishes.

## Supporting Information

**Figure S1: Stable isotope analysis of the trophic samples from the benthic samples**

**Figure S2: Stable isotope analysis of the trophic samples from the stomach contents**

**Figure S3: Trophic positions comparisons based on stomach content baseline**

**Table S1: sample list overview of muscle samples isotope values and details about sampling**

**Table S2: sample list overview of trophic samples**

**Table S3: results of single-factor ANOVAs and Tukey-Kramer test (if applicable) for species with transition season samples testing the factor of locality**

## Acknowledgements

We deeply thank Monika Kłodawska, Dima Omelchenko and Hassan Bassirou for the field assistance as well as their valuable support as colleagues. We are very thankful to the local fisherman Raphael Abanda without whom no successful fishing would have been possible and whose knowledge of the area was indispensable. We further thank our field guides Jean-Claude Remix and Gregoire Kayoum from the “Institut de Recherche pour le Développement (IRD)”, as well as our hosts in the Batchenga village. We sincerely express gratitude to the local people of the Batchenga community to allow us to fish in their territory, the Nachtigal HydroPower Company (NHPC) for the support to sample in the Sanaga and the Mpem and Djim National Park rivers. We thank all the students and people from the Institute for Environmental Studies and the Department of Botany of Charles University for sharing space and giving us the possibility to work there and offering kind advice and friendliness. A special thanks goes to Marc Schmid for providing the map in Figure 1.

## Author contribution statement

Conceptualisation: ZM, ARBN. Developing methods: ZM, ARBN, PH, JK Conducting the research: ZM, AI, SDN, PH, GMS. Data analysis: PH, JK, GMS. Data interpretation: ZM, SDN, GMS. Preparation figures and tables: GMS. Writing – initial draft - GMS and ZM, Writing – final revision and approval – all authors.

## Data Availability statement

COI sequences used for the species phylogeny are available on GenBank for *Marcusenius mento* with accession numbers: HM882757 and HM882727, and (will be available) for the other species with the accession numbers (insert). Raw data of stable isotope analysis of the mass spectrometer for muscle samples are imbedded in sample overview table in Table S1 in Supporting information and for trophic samples in Table S2.

