## Supplemental Figures S1-3 for "Trophic ecology of the African riverine elephant fishes (Mormyridae)"

Supplementary Material for:

Gina Maria Sommer, Samuel Didier Njom, Adrian Indermaur, Arnold Roger Bitja Nyom, Petra Horká, Jaroslav Kukla, Zuzana Musilová - **Trophic niche differentiation in the African elephant fish community (Mormyridae) from the Sanaga river in Cameroon**

**Figure S1: Stable isotope analysis of the trophic samples from the benthic samples**

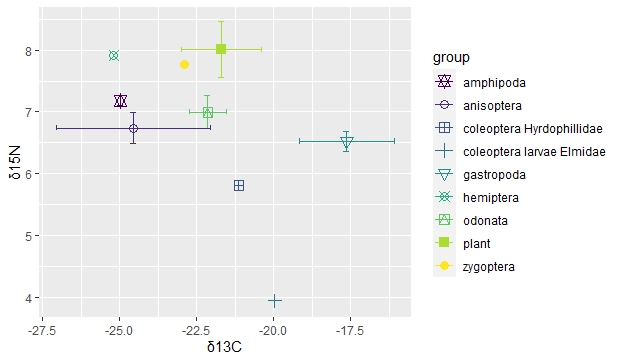

**Figure S2: Stable isotope analysis of the trophic samples from the stomach contents**

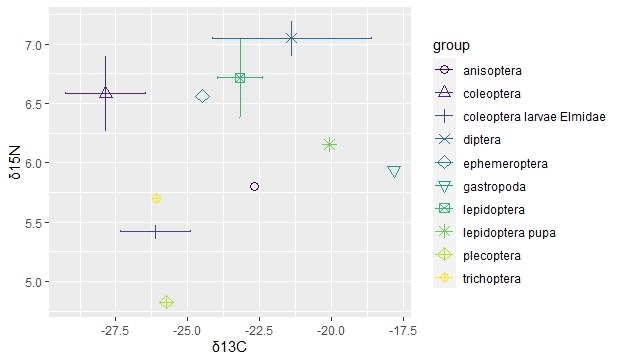

**Figure S3: Trophic positions comparisons based on stomach content baseline**

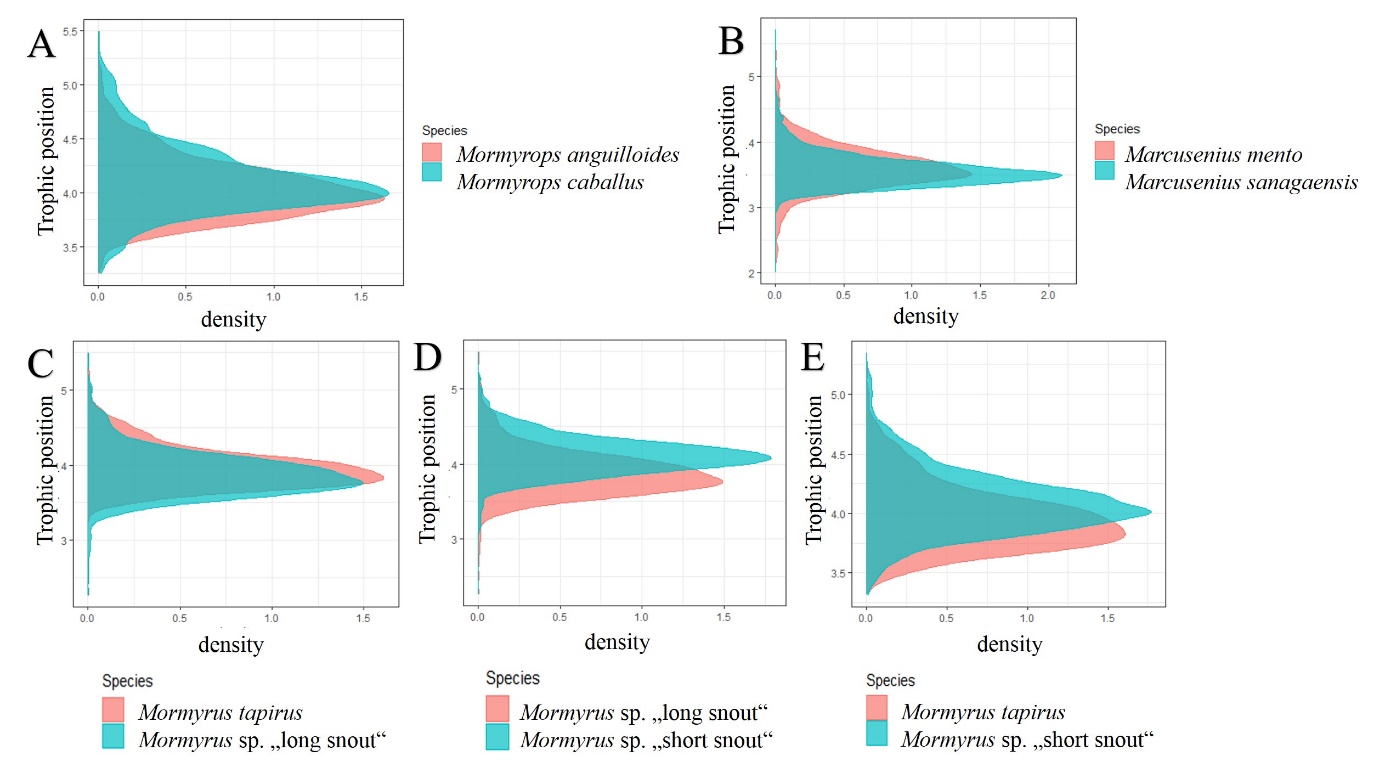

**Table S1: sample list overview of muscle samples isotope values and details about sampling**

**Table S2: sample list overview of trophic samples**

**Table S3: results of single-factor ANOVAs and Tukey-Kramer test (if applicable) for species with transition season samples testing the factor of locality**

Provided together in a separate pdf file
