## Supplemental Tables S1-S3 for "Trophic ecology of the African riverine elephant fishes (Mormyridae)"

| **code** | ***species*** | **d15N** | **d13C** | **Season** | **date** | **month** | **year** | **location** |
|---|---|---|---|---|---|---|---|---|
| **62D3** | *Campylomormyrus phantasticus* | 12.236 | -23.124 | dry | 8 | March | 2017 | Sanaga, Nachtigal falls |
| **62D9** | *Campylomormyrus phantasticus* | 12.253 | -22.924 | dry | 8 | March | 2017 | Sanaga, Nachtigal falls |
| **62F6** | *Campylomormyrus phantasticus* | 12.016 | -22.753 | dry | 10 | March | 2017 | Sanaga, Nachtigal falls |
| **62C2** | *Campylomormyrus phantasticus* | 12.262 | -22.922 | dry | 8 | March | 2017 | Sanaga, Nachtigal falls |
| **16338** | *Campylomormyrus phantasticus* | 12.505 | -23.715 | wet | 1 | November | 2018 | Sanaga, Nachtigal falls |
| **16339** | *Campylomormyrus phantasticus* | 11.942 | -23.265 | wet | 1 | November | 2018 | Sanaga, Nachtigal falls |
| **62G1** | *Campylomormyrus phantasticus* | 12.278 | -23.495 | dry | 11 | March | 2017 | Sanaga, Nachtigal falls |
| **101B5** | *Campylomormyrus phantasticus* | 12.619 | -24.207 | dry | 19 | February | 2018 | Sanaga, Nachtigal falls |
| **101D7** | *Campylomormyrus phantasticus* | 12.430 | -23.917 | dry | 19 | February | 2018 | Sanaga, Nachtigal falls |
| **101E2** | *Campylomormyrus phantasticus* | 12.195 | -23.924 | dry | 20 | February | 2018 | Sanaga, Nachtigal falls, upstream |
| **101E3** | *Campylomormyrus phantasticus* | 12.017 | -25.436 | dry | 20 | February | 2018 | Sanaga, Nachtigal falls, upstream |
| **101E4** | *Campylomormyrus phantasticus* | 12.333 | -23.535 | dry | 20 | February | 2018 | Sanaga, Nachtigal falls, upstream |
| **101G6** | *Campylomormyrus phantasticus* | 12.170 | -24.302 | dry | 21 | February | 2018 | Avo'o |
| **15503** | *Campylomormyrus phantasticus* | 12.040 | -22.252 | transition | 11 | July | 2018 | Sanaga |
| **15512** | *Campylomormyrus phantasticus* | 12.184 | -22.964 | transition | 11 | July | 2018 | Sanaga |
| **15519** | *Campylomormyrus phantasticus* | 12.141 | -22.568 | transition | 11 | July | 2018 | Sanaga |
| **15520** | *Campylomormyrus phantasticus* | 12.229 | -22.132 | transition | 11 | July | 2018 | Sanaga |
| **15558** | *Campylomormyrus phantasticus* | 12.261 | -23.389 | wet | 5 | September | 2018 | Sanaga, Nachtigal falls |
| **15601** | *Campylomormyrus phantasticus* | 12.428 | -23.519 | wet | 5 | September | 2018 | Sanaga, Nachtigal falls |
| **14345** | *Campylomormyrus phantasticus* | 12.139 | -23.152 | dry | 21 | March | 2018 | Sanaga, Nachtigal falls, upstream |
| **14363** | *Campylomormyrus phantasticus* | 12.138 | -24.563 | dry | 21 | March | 2018 | Sanaga, Nachtigal falls, upstream |
| **14364** | *Campylomormyrus phantasticus* | 12.092 | -24.557 | dry | 21 | March | 2018 | Sanaga, Nachtigal falls, upstream |
| **101G8** | *Campylomormyrus phantasticus* | 12.059 | -24.449 | dry | 21 | February | 2018 | Avo'o |
| **14183** | *Campylomormyrus phantasticus* | 12.184 | -24.157 | dry | 20 | February | 2018 | Sanaga, Nachtigal falls, upstream |
| **14184** | *Campylomormyrus phantasticus* | 12.164 | -23.947 | dry | 20 | February | 2018 | Sanaga, Nachtigal falls, upstream |
| **14401** | *Campylomormyrus phantasticus* | 12.161 | -23.899 | dry | 22 | March | 2018 | Sanaga, Nachtigal falls, upstream |

| code | *species* | d15N | d13C | Season | date | month | year | location |
|---|---|---|---|---|---|---|---|---|
| **15711** | *Campylomormyrus phantasticus* | 12.126 | -24.339 | wet | 8 | September | 2018 | Sanaga, Nachtigal falls, upstream |
| **15712** | *Campylomormyrus phantasticus* | 12.150 | -23.381 | wet | 8 | September | 2018 | Sanaga, Nachtigal falls, upstream |
| **15713** | *Campylomormyrus phantasticus* | 12.189 | -23.386 | wet | 8 | September | 2018 | Sanaga, Nachtigal falls, upstream |
| **62A2** | *Campylomormyrus phantasticus* | 11.871 | -22.532 | dry | 7 | March | 2017 | Sanaga, Nachtigal falls |
| **62C5** | *Campylomormyrus phantasticus* | 11.779 | -22.193 | dry | 8 | March | 2017 | Sanaga, Nachtigal falls |
| **62C6** | *Campylomormyrus phantasticus* | 11.999 | -22.887 | dry | 8 | March | 2017 | Sanaga, Nachtigal falls |
| **62D7** | *Hippopotamyrus castor* | 11.335 | -23.877 | dry | 9 | March | 2017 | Sanaga, Nachtigal falls |
| **62A4** | *Hippopotamyrus castor* | 12.259 | -23.487 | dry | 7 | March | 2017 | Sanaga, Nachtigal falls |
| **62A5** | *Hippopotamyrus castor* | 12.055 | -23.541 | dry | 7 | March | 2017 | Sanaga, Nachtigal falls |
| **101D5** | *Hippopotamyrus castor* | 12.610 | -24.161 | dry | 19 | February | 2018 | Sanaga, Nachtigal falls |
| **101D9** | *Hippopotamyrus castor* | 12.165 | -24.302 | dry | 20 | February | 2018 | Sanaga, Nachtigal falls, upstream |
| **16340** | *Hippopotamyrus castor* | 12.755 | -24.085 | wet | 1 | November | 2018 | Sanaga, Nachtigal falls |
| **16341** | *Hippopotamyrus castor* | 12.315 | -23.269 | wet | 1 | November | 2018 | Sanaga, Nachtigal falls |
| **101E1** | *Hippopotamyrus castor* | 11.561 | -19.593 | dry | 20 | February | 2018 | Sanaga, Nachtigal falls,upstream |
| **101G5** | *Hippopotamyrus castor* | 12.444 | -24.063 | dry | 21 | February | 2018 | Avo'o |
| **15526** | *Hippopotamyrus castor* | 11.485 | -24.020 | transition | 11 | July | 2018 | Sanaga, Nachtigal falls |
| **13582** | *Hippopotamyrus castor* | 10.367 | -28.910 | dry | 12 | February | 2018 | Stream park (Djim river) |
| **15071** | *Hippopotamyrus castor* | 11.183 | -29.208 | transition | 29 | May | 2018 | Stream park |
| **12402** | *Hippopotamyrus castor* | 11.635 | -23.155 | wet | 8 | November | 2017 | Stream park (Mey river) |
| **14344** | *Hippopotamyrus castor* | 11.497 | -23.007 | dry | 21 | March | 2018 | Sanaga, Nachtigal falls,upstream |
| **15667** | *Hippopotamyrus castor* | 11.842 | -21.219 | wet | 7 | September | 2018 | Sanaga, Nachtigal falls,upstream |
| **15698** | *Hippopotamyrus castor* | 11.820 | -23.080 | wet | 7 | September | 2018 | Sanaga, Nachtigal falls,upstream |
| **12464** | *Hippopotamyrus castor* | 12.484 | -23.733 | wet | 9 | November | 2017 | Sanaga, Nachtigal falls,upstream |
| **14189** | *Hippopotamyrus castor* | 11.458 | -19.511 | dry | 20 | February | 2018 | Sanaga, Nachtigal falls,upstream |
| **12313** | *Marcusenius mento* | 11.386 | -25.798 | wet | 25 | August | 2017 | Stream park (Mpem river) |
| **14819** | *Marcusenius mento* | 10.313 | -31.395 | transition | 25 | May | 2018 | Stream park (Mpem river) |
| **14833** | *Marcusenius mento* | 11.227 | -29.672 | transition | 26 | May | 2018 | Stream park (Mpem river) |

| **code** | ***species*** | **d15N** | **d13C** | **Season** | **date** | **month** | **year** | **location** |
|---|---|---|---|---|---|---|---|---|
| **14860** | *Marcusenius mento* | 10.308 | -30.384 | transition | 26 | May | 2018 | Stream park (Mpem river) |
| **12078** | *Marcusenius sanagaensis* | 12.381 | -24.519 | wet | 16 | September | 2017 | Stream san (Asamba river) |
| **12084** | *Marcusenius sanagaensis* | 10.340 | -27.333 | wet | 16 | Oktober | 2017 | Stream san (Asamba river) |
| **14452** | *Marcusenius sanagaensis* | 11.061 | -27.501 | dry | 23 | March | 2018 | Stream san (Asamba river) |
| **12137** | *Marcusenius sanagaensis* | 10.433 | -28.338 | wet | 17 | August | 2017 | Stream san (Nia river) |
| **62F9** | *Marcusenius sanagaensis* | 10.907 | -16.514 | dry | 11 | March | 2017 | Sanaga, Nachtigal falls |
| **62D4** | *Marcusenius sanagaensis* | 11.922 | -20.360 | dry | 8 | March | 2017 | Sanaga, Nachtigal falls |
| **12085** | *Marcusenius sanagaensis* | 10.327 | -27.288 | wet | 16 | August | 2017 | Stream san (Asamba river) |
| **12086** | *Marcusenius sanagaensis* | 10.227 | -26.837 | wet | 16 | August | 2017 | Stream san (Asamba river) |
| **12469** | *Marcusenius sanagaensis* | 10.168 | -26.533 | wet | 10 | November | 2017 | Stream san (Asamba river) |
| **12477** | *Marcusenius sanagaensis* | 10.214 | -27.633 | wet | 10 | November | 2017 | Stream san (Asamba river) |
| **14497** | *Marcusenius sanagaensis* | 10.523 | -27.795 | dry | 23 | March | 2018 | Stream san (Asamba river) |
| **14498** | *Marcusenius sanagaensis* | 10.587 | -27.683 | dry | 23 | March | 2018 | Stream san (Asamba river) |
| **14499** | *Marcusenius sanagaensis* | 10.876 | -26.652 | dry | 23 | March | 2018 | Stream san (Asamba river) |
| **15212** | *Marcusenius sanagaensis* | 10.433 | -28.107 | transition | 7 | June | 2018 | Stream san (Asamba river) |
| **15216** | *Marcusenius sanagaensis* | 10.145 | -28.087 | transition | 7 | June | 2018 | Stream san (Asamba river) |
| **15536** | *Marcusenius sanagaensis* | 10.757 | -27.092 | transition | 12 | July | 2018 | Stream san (Asamba river) |
| **15788** | *Marcusenius sanagaensis* | 10.503 | -27.383 | wet | 9 | September | 2018 | Stream san (Asamba river) |
| **12187** | *Marcusenius sanagaensis* | 10.407 | -27.290 | wet | 19 | August | 2017 | Stream san (Banga river) |
| **14586** | *Marcusenius sanagaensis* | 9.416 | -23.757 | dry | 25 | March | 2018 | Stream san (Banga river) |
| **14587** | *Marcusenius sanagaensis* | 9.382 | -30.131 | dry | 25 | March | 2018 | Stream san (Banga river) |
| **13680** | *Marcusenius sanagaensis* | 9.954 | -28.176 | dry | 13 | February | 2018 | Stream park (Djim river) |
| **13681** | *Marcusenius sanagaensis* | 10.952 | -27.767 | dry | 13 | February | 2018 | Stream park (Djim river) |
| **13682** | *Marcusenius sanagaensis* | 10.641 | -29.134 | dry | 13 | February | 2018 | Stream park (Djim river) |
| **13683** | *Marcusenius sanagaensis* | 10.289 | -27.392 | dry | 13 | February | 2018 | Stream park (Djim river) |
| **13684** | *Marcusenius sanagaensis* | 10.356 | -26.553 | dry | 13 | February | 2018 | Stream park (Djim river) |
| **15342** | *Marcusenius sanagaensis* | 8.787 | -16.720 | transition | 9 | June | 2018 | Stream san (Mekono river) |

| code | *species* | d15N | d13C | Season | date | month | year | location |
|---|---|---|---|---|---|---|---|---|
| **15343** | *Marcusenius sanagaensis* | 9.053 | -16.505 | transition | 9 | June | 2018 | Stream san (Mekono river) |
| **15344** | *Marcusenius sanagaensis* | 9.023 | -15.979 | transition | 9 | June | 2018 | Stream san (Mekono river) |
| **15346** | *Marcusenius sanagaensis* | 8.731 | -17.838 | transition | 9 | June | 2018 | Stream san (Mekono river) |
| **15070** | *Marcusenius sanagaensis* | 10.605 | -27.595 | transition | 29 | May | 2018 | Stream park (Mey river) |
| **12594** | *Marcusenius sanagaensis* | 10.706 | -29.910 | wet | 21 | November | 2017 | Stream park (Mpem river) |
| **12619** | *Marcusenius sanagaensis* | 11.551 | -26.974 | wet | 21 | November | 2017 | Stream park (Mpem river) |
| **12620** | *Marcusenius sanagaensis* | 11.201 | -26.750 | wet | 21 | November | 2017 | Stream park (Mpem river) |
| **14831** | *Marcusenius sanagaensis* | 9.962 | -27.503 | transition | 26 | May | 2018 | Stream park (Mpem river) |
| **14832** | *Marcusenius sanagaensis* | 10.906 | -27.292 | transition | 26 | May | 2018 | Stream park (Mpem river) |
| **12114** | *Marcusenius sanagaensis* | 10.763 | -25.985 | wet | 17 | August | 2017 | Stream san (Nia river) |
| **12136** | *Marcusenius sanagaensis* | 10.156 | -27.251 | wet | 17 | August | 2017 | Stream san (Nia river) |
| **14548** | *Marcusenius sanagaensis* | 10.650 | -28.024 | dry | 24 | March | 2018 | Stream san (Nia river) |
| **15920** | *Marcusenius sanagaensis* | 10.827 | -25.661 | wet | 14 | September | 2018 | Stream san (Nia river) |
| **15921** | *Marcusenius sanagaensis* | 10.511 | -26.193 | wet | 14 | September | 2018 | Stream san (Nia river) |
| **12556** | *Marcusenius sanagaensis* | 9.701 | -27.417 | wet | 15 | November | 2017 | Stream san (Seele river) |
| **14690** | *Marcusenius sanagaensis* | 10.782 | -29.751 | dry | 27 | March | 2018 | Stream san (Sele river) |
| **16136** | *Marcusenius sanagaensis* | 10.699 | -27.806 | wet | 16 | September | 2018 | Stream san (Tede river) |
| **62A8** | *Mormyrops anguilloides* | 11.624 | -24.941 | dry | 7 | March | 2017 | Sanaga, Nachtigal falls |
| **62AB3** | *Mormyrops anguilloides* | 12.918 | -22.165 | dry | 7 | March | 2017 | Sanaga, Nachtigal falls |
| **62B4** | *Mormyrops anguilloides* | 12.253 | -24.423 | dry | 7 | March | 2017 | Sanaga, Nachtigal falls |
| **62C1** | *Mormyrops anguilloides* | 12.985 | -21.488 | dry | 7 | March | 2017 | Sanaga, Nachtigal falls |
| **15219** | *Mormyrops anguilloides* | 12.739 | -26.394 | transition | 7 | June | 2018 | Stream san (Asamba river) |
| **12035** | *Mormyrops anguilloides* | 12.504 | -23.730 | wet | 14 | August | 2017 | Stream san (Avo’o river) |
| **13943** | *Mormyrops anguilloides* | 12.556 | -34.964 | dry | 18 | February | 2018 | Sanaga, Nachtigal falls |
| **101A8** | *Mormyrops anguilloides* | 12.163 | -25.657 | dry | 18 | February | 2018 | Sanaga, Nachtigal falls |
| **16267** | *Mormyrops anguilloides* | 12.310 | -23.132 | wet | 29 | October | 2018 | Sanaga, Nachtigal falls |
| **16290** | *Mormyrops anguilloides* | 13.024 | -22.971 | wet | 30 | October | 2018 | Sanaga, Nachtigal falls |

| **code** | ***species*** | **d15N** | **d13C** | **Season** | **date** | **month** | **year** | **location** |
|---|---|---|---|---|---|---|---|---|
| **16303** | *Mormyrops anguilloides* | 12.114 | -24.866 | wet | 31 | October | 2018 | Sanaga, Nachtigal falls |
| **16304** | *Mormyrops anguilloides* | 12.352 | -24.324 | wet | 31 | October | 2018 | Sanaga, Nachtigal falls |
| **16305** | *Mormyrops anguilloides* | 11.134 | -26.195 | wet | 31 | October | 2018 | Sanaga, Nachtigal falls |
| **16311** | *Mormyrops anguilloides* | 11.632 | -28.465 | wet | 31 | October | 2018 | Sanaga, Nachtigal falls |
| **16312** | *Mormyrops anguilloides* | 10.914 | -29.213 | wet | 31 | October | 2018 | Sanaga, Nachtigal falls |
| **16313** | *Mormyrops anguilloides* | 12.788 | -23.492 | wet | 31 | October | 2018 | Sanaga, Nachtigal falls |
| **16314** | *Mormyrops anguilloides* | 12.622 | -24.075 | wet | 1 | November | 2018 | Stream san (Wala river) |
| **16326** | *Mormyrops anguilloides* | 12.185 | -22.219 | wet | 1 | November | 2018 | Stream san (Wala river) |
| **16332** | *Mormyrops anguilloides* | 11.019 | -26.016 | wet | 1 | November | 2018 | Sanaga, Nachtigal falls |
| **16333** | *Mormyrops anguilloides* | 12.742 | -23.729 | wet | 1 | November | 2018 | Sanaga, Nachtigal falls |
| **101C6** | *Mormyrops anguilloides* | 13.124 | -24.278 | dry | 19 | February | 2018 | Sanaga, Nachtigal falls |
| **13446** | *Mormyrops anguilloides* | 11.308 | -28.121 | dry | 12 | February | 2018 | Stream park (Djim river) |
| **12408** | *Mormyrops anguilloides* | 11.710 | -26.588 | wet | 8 | November | 2017 | Sanaga, Nachtigal falls,upstream |
| **14391** | *Mormyrops anguilloides* | 11.508 | -30.994 | dry | 21 | March | 2018 | Sanaga, Nachtigal falls,upstream |
| **12105** | *Mormyrops anguilloides* | 12.093 | -25.032 | wet | 17 | August | 2017 | Stream san (Nia river) |
| **12555** | *Mormyrops anguilloides* | 11.726 | -28.883 | wet | 15 | November | 2017 | Stream san (Sele river) |
| **62D1** | *Mormyrops anguilloides* | 12.730 | -22.191 | dry | 8 | March | 2017 | Sanaga, Nachtigal falls |
| **62D2** | *Mormyrops anguilloides* | 12.090 | -22.875 | dry | 8 | March | 2017 | Sanaga, Nachtigal falls |
| **62E2** | *Mormyrops anguilloides* | 11.943 | -24.203 | dry | 9 | March | 2017 | Sanaga, Nachtigal falls |
| **62F1** | *Mormyrops anguilloides* | 11.792 | -23.266 | dry | 9 | March | 2017 | Sanaga, Nachtigal falls |
| **62G3** | *Mormyrops anguilloides* | 11.753 | -27.546 | dry | 11 | March | 2017 | Sanaga, Nachtigal falls |
| **12034** | *Mormyrops caballus* | 12.715 | -22.931 | wet | 14 | August | 2017 | Stream san (Avo'o river) |
| **16291** | *Mormyrops caballus* | 12.873 | -21.914 | wet | 30 | October | 2018 | Sanaga, Nachtigal falls |
| **62F3** | *Mormyrops caballus* | 12.328 | -22.292 | dry | 10 | March | 2017 | Asamba river |
| **62G4** | *Mormyrops caballus* | 12.347 | -22.700 | dry | 11 | March | 2017 | Sanaga, Nachtigal falls |
| **101A9** | *Mormyrops caballus* | 12.460 | -22.254 | dry | 18 | February | 2018 | Sanaga, Nachtigal falls |
| **16334** | *Mormyrops caballus* | 12.489 | -23.662 | wet | 1 | November | 2018 | Sanaga, Nachtigal falls |

| code | *species* | d15N | d13C | Season | date | month | year | location |
|---|---|---|---|---|---|---|---|---|
| **16342** | *Mormyrops caballus* | 12.126 | -23.492 | wet | 2 | November | 2018 | Stream san (Wala river) |
| **101B4** | *Mormyrops caballus* | 12.302 | -24.865 | dry | 19 | February | 2018 | Sanaga, Nachtigal falls |
| **101C7** | *Mormyrops caballus* | 12.483 | -22.575 | dry | 19 | February | 2018 | Sanaga, Nachtigal falls |
| **101C4** | *Mormyrops caballus* | 11.689 | -24.538 | dry | 19 | February | 2018 | Sanaga, Nachtigal falls |
| **101F7** | *Mormyrops caballus* | 12.555 | -22.037 | dry | 21 | February | 2018 | Avo'o |
| **101F8** | *Mormyrops caballus* | 12.292 | -22.219 | dry | 21 | February | 2018 | Avo'o |
| **14005** | *Mormyrops caballus* | 12.338 | -22.291 | dry | 19 | February | 2018 | Sanaga, Nachtigal falls |
| **14431** | *Mormyrops caballus* | 12.635 | -21.889 | dry | 22 | March | 2018 | Sanaga, Nachtigal falls, upstream |
| **14443** | *Mormyrops caballus* | 12.453 | -22.631 | dry | 22 | March | 2018 | Sanaga, Nachtigal falls, upstream |
| **14675** | *Mormyrops caballus* | 11.857 | -29.457 | dry | 27 | March | 2018 | Stream san (Sele river) |
| **14676** | *Mormyrops caballus* | 11.890 | -29.351 | dry | 27 | March | 2018 | Stream san (Sele river) |
| **15981** | *Mormyrops caballus* | 11.826 | -30.549 | wet | 15 | September | 2018 | Stream san (Sele river) |
| **14207** | *Mormyrops caballus* | 12.460 | -22.049 | dry | 21 | February | 2018 | Sanaga, Nachtigal falls, upstream |
| **14218** | *Mormyrops caballus* | 12.207 | -22.233 | dry | 21 | February | 2018 | Sanaga, Nachtigal falls, upstream |
| **62B8** | *Mormyrops caballus* | 12.326 | -23.362 | dry | 7 | March | 2017 | Sanaga, Nachtigal falls |
| **62C8** | *Mormyrus tapirus* | 12.150 | -24.227 | dry | 8 | March | 2017 | Sanaga, Nachtigal falls |
| **62C9** | *Mormyrus tapirus* | 12.493 | -24.722 | dry | 8 | March | 2017 | Sanaga, Nachtigal falls |
| **62D5** | *Mormyrus tapirus* | 11.352 | -26.149 | dry | 8 | March | 2017 | Sanaga, Nachtigal falls |
| **62D6** | *Mormyrus tapirus* | 11.976 | -23.856 | dry | 8 | March | 2017 | Sanaga, Nachtigal falls |
| **62D8** | *Mormyrus tapirus* | 9.520 | -26.765 | dry | 9 | March | 2017 | Sanaga, Nachtigal falls |
| **62E5** | *Mormyrus tapirus* | 11.953 | -24.133 | dry | 9 | March | 2017 | Sanaga, Nachtigal falls |
| **62E6** | *Mormyrus tapirus* | 12.107 | -23.693 | dry | 9 | March | 2017 | Sanaga, Nachtigal falls |
| **62E8** | *Mormyrus tapirus* | 11.059 | -24.568 | dry | 9 | March | 2017 | Sanaga, Nachtigal falls |
| **62E7** | *Mormyrus tapirus* | 11.638 | -24.148 | dry | 9 | March | 2017 | Sanaga, Nachtigal falls |
| **62A1** | *Mormyrus tapirus* | 10.851 | -24.324 | dry | 7 | March | 2017 | Sanaga, Nachtigal falls |
| **62A9** | *Mormyrus tapirus* | 10.830 | -24.410 | dry | 7 | March | 2017 | Sanaga, Nachtigal falls |
| **62B9** | *Mormyrus tapirus* | 11.605 | -24.940 | dry | 7 | March | 2017 | Sanaga, Nachtigal falls |

| **code** | ***species*** | **d15N** | **d13C** | **Season** | **date** | **month** | **year** | **location** |
|---|---|---|---|---|---|---|---|---|
| **62C3** | *Mormyrus tapirus* | 13.474 | -25.555 | dry | 8 | March | 2017 | Sanaga, Nachtigal falls |
| **62G2** | *Mormyrus tapirus* | 11.871 | -21.922 | dry | 11 | March | 2017 | Sanaga, Nachtigal falls |
| **101C8** | *Mormyrus tapirus* | 12.309 | -20.441 | dry | 19 | February | 2018 | Sanaga, Nachtigal falls |
| **101 E8** | *Mormyrus tapirus* | 12.322 | -23.204 | dry | 20 | February | 2018 | Sanaga, Nachtigal falls,upstream |
| **101A7** | *Mormyrus tapirus* | 12.447 | -24.355 | dry | 18 | February | 2018 | Sanaga, Nachtigal falls |
| **101B1** | *Mormyrus tapirus* | 11.688 | -24.368 | dry | 18 | February | 2018 | Sanaga, Nachtigal falls |
| **101B2** | *Mormyrus tapirus* | 11.385 | -23.723 | dry | 18 | February | 2018 | Sanaga, Nachtigal falls |
| **101B3** | *Mormyrus tapirus* | 10.726 | -27.153 | dry | 18 | February | 2018 | Sanaga, Nachtigal falls |
| **101B6** | *Mormyrus tapirus* | 11.576 | -25.294 | dry | 19 | February | 2018 | Sanaga, Nachtigal falls |
| **101B8** | *Mormyrus tapirus* | 11.281 | -25.500 | dry | 19 | February | 2018 | Sanaga, Nachtigal falls |
| **101C1** | *Mormyrus tapirus* | 12.833 | -21.620 | dry | 19 | February | 2018 | Sanaga, Nachtigal falls |
| **101B9** | *Mormyrus tapirus* | 11.919 | -21.148 | dry | 19 | February | 2018 | Sanaga, Nachtigal falls |
| **101C3** | *Mormyrus tapirus* | 12.175 | -23.541 | dry | 19 | February | 2018 | Sanaga, Nachtigal falls |
| **101C9** | *Mormyrus tapirus* | 11.904 | -24.412 | dry | 19 | February | 2018 | Sanaga, Nachtigal falls |
| **16281** | *Mormyrus tapirus* | 10.870 | -23.804 | wet | 30 | October | 2018 | Sanaga, Nachtigal falls |
| **16293** | *Mormyrus tapirus* | 12.684 | -23.020 | wet | 30 | October | 2018 | Sanaga, Nachtigal falls |
| **15523** | *Mormyrus tapirus* | 12.066 | -23.650 | transition | 11 | July | 2018 | Sanaga, Nachtigal falls, upstream |
| **15524** | *Mormyrus tapirus* | 9.359 | -28.020 | transition | 11 | July | 2018 | Sanaga, Nachtigal falls, upstream |
| **14203** | *Mormyrus tapirus* | 12.031 | -24.422 | dry | 21 | February | 2018 | Sanaga, Nachtigal falls, upstream |
| **14204** | *Mormyrus tapirus* | 12.017 | -22.717 | dry | 21 | February | 2018 | Sanaga, Nachtigal falls, upstream |
| **14205** | *Mormyrus tapirus* | 11.951 | -20.869 | dry | 21 | February | 2018 | Sanaga, Nachtigal falls, upstream |
| **12361** | *Mormyrus tapirus* | 12.000 | -22.953 | wet | 6 | November | 2017 | Stream san (Wala river) |
| **14187** | *Mormyrus tapirus* | 11.83 | -24.065 | dry | 20 | February | 2018 | Sanaga, Nachtigal falls, upstream |
| **14418** | *Mormyrus tapirus* | 13.454 | -22.491 | dry | 22 | March | 2018 | Sanaga, Nachtigal falls, upstream |
| **16337** | *Mormyrus tapirus* | 11.617 | -22.912 | wet | 1 | November | 2018 | Sanaga, Nchtigal falls |
| **16347** | *Mormyrus tapirus* | 11.532 | -22.833 | wet | 2 | November | 2018 | Stream san (Wala river) |
| **16348** | *Mormyrus tapirus* | 11.750 | -21.858 | wet | 2 | November | 2018 | Stream san (Wala river) |

| **code** | ***species*** | **d15N** | **d13C** | **Season** | **date** | **month** | **year** | **location** |
|---|---|---|---|---|---|---|---|---|
| **16315** | *Mormyrus tapirus* | 11.763 | -24.233 | wet | 1 | November | 2018 | Stream san (Wala river) |
| **16321** | *Mormyrus tapirus* | 11.671 | -23.794 | wet | 1 | November | 2018 | Stream san (Wala river) |
| **16323** | *Mormyrus tapirus* | 12.198 | -23.063 | wet | 1 | November | 2018 | Stream san (Wala river) |
| **16324** | *Mormyrus tapirus* | 12.753 | -23.218 | wet | 1 | November | 2018 | Stream san (Wala river) |
| **16335** | *Mormyrus tapirus* | 12.224 | -23.426 | wet | 1 | November | 2018 | Sanaga, Nachtigal falls |
| **16336** | *Mormyrus tapirus* | 12.289 | -23.361 | wet | 1 | November | 2018 | Sanaga, Nachtigal falls |
| **16346** | *Mormyrus tapirus* | 11.865 | -23.129 | wet | 2 | November | 2018 | Stream san (Wala river) |
| **15037** | *Mormyrus tapirus* | 10.902 | -28.512 | transition | 29 | May | 2018 | Stream park (Mpem river) |
| **15045** | *Mormyrus tapirus* | 9.859 | -30.128 | transition | 29 | May | 2018 | Stream park (Mpem river) |
| **14825** | *Mormyrus tapirus* | 9.888 | -30.049 | transition | 26 | May | 2018 | Stream park (Mpem river) |
| **14835** | *Mormyrus tapirus* | 11.454 | -27.864 | transition | 26 | May | 2018 | Stream park (Mpem river) |
| **14346** | *Mormyrus tapirus* | 11.305 | -26.256 | dry | 21 | March | 2018 | Sanaga, Nachtigal falls, upstream |
| **12360** | *Mormyrus tapirus* | 11.775 | -22.858 | wet | 6 | November | 2017 | Stream san (Wala river) |
| **62A3** | *Mormyrus* sp. "long snout" | 11.490 | -23.122 | dry | 7 | March | 2017 | Sanaga, Nachtigal falls |
| **62A6** | *Mormyrus* sp. "long snout" | 11.443 | -23.367 | dry | 7 | March | 2017 | Sanaga, Nachtigal falls |
| **62A7** | *Mormyrus* sp. "long snout" | 11.329 | -22.697 | dry | 7 | March | 2017 | Sanaga, Nachtigal falls |
| **12160** | *Mormyrus* sp. "short snout" | 12.558 | -20.070 | wet | 18 | August | 2017 | Sanaga, Nachtigal falls, upstream |
| **12445** | *Mormyrus* sp. "short snout" | 12.963 | -21.937 | wet | 9 | November | 2017 | Sanaga, Nachtigal falls, upstream |
| **12446** | *Mormyrus* sp. "short snout" | 12.431 | -20.701 | wet | 9 | November | 2017 | Sanaga, Nachtigal falls, upstream |
| **14181** | *Mormyrus* sp. "short snout" | 12.673 | -22.421 | dry | 20 | February | 2018 | Sanaga, Nachtigal falls, upstream |
| **14185** | *Mormyrus* sp. "short snout" | 12.326 | -20.721 | dry | 20 | February | 2018 | Sanaga, Nachtigal falls, upstream |
| **14206** | *Mormyrus* sp. "short snout" | 11.981 | -22.450 | dry | 21 | February | 2018 | Sanaga, Nachtigal falls, upstream |
| **15730** | *Mormyrus* sp. "short snout" | 12.435 | -21.635 | wet | 8 | September | 2018 | Sanaga, Nachtigal falls, upstream |
| **15734** | *Mormyrus* sp. "short snout" | 12.599 | -20.167 | wet | 8 | September | 2018 | Sanaga, Nachtigal falls, upstream |
| **15699** | *Mormyrus* sp. "short snout" | 12.134 | -19.847 | wet | 7 | September | 2018 | Sanaga, Nachtigal falls, upstream |
| **62C7** | *Mormyrus* sp. "short snout" | 11.750 | -20.700 | dry | 7 | March | 2017 | Sanaga, Nachtigal falls |
| **62B1** | *Mormyrus* sp. "short snout" | 11.960 | -20.014 | dry | 7 | March | 2017 | Sanaga, Nachtigal falls |

| code | *species* | d15N | d13C | Season | date | month | year | location |
|---|---|---|---|---|---|---|---|---|
| **62B2** | *Mormyrus* sp. "short snout" | 12.097 | -18.485 | dry | 7 | March | 2017 | Sanaga, Nachtigal falls |
| **62B5** | *Mormyrus* sp. "short snout" | 11.875 | -20.895 | dry | 7 | March | 2017 | Sanaga, Nachtigal falls |
| **62B6** | *Mormyrus* sp. "short snout" | 12.021 | -22.138 | dry | 7 | March | 2017 | Sanaga, Nachtigal falls |
| **62B7** | *Mormyrus* sp. "short snout" | 12.458 | -21.862 | dry | 7 | March | 2017 | Sanaga, Nachtigal falls |
| **62C4** | *Mormyrus* sp. "short snout" | 12.490 | -22.204 | dry | 8 | March | 2017 | Sanaga, Nachtigal falls |
| **62F8** | *Mormyrus* sp. "short snout" | 12.163 | -22.555 | dry | 11 | March | 2017 | Sanaga, Nachtigal falls |
| **62G5** | *Mormyrus* sp. "short snout" | 11.248 | -26.525 | dry | 11 | March | 2017 | Sanaga, Nachtigal falls |
| **101A6** | *Mormyrus* sp. "short snout" | 11.876 | -18.652 | dry | 18 | February | 2018 | Sanaga, Nachtigal falls |
| **101B7** | *Mormyrus* sp. "short snout" | 12.638 | -19.068 | dry | 19 | February | 2018 | Sanaga, Nachtigal falls |
| **101C2** | *Mormyrus* sp. "short snout" | 12.427 | -19.405 | dry | 19 | February | 2018 | Sanaga, Nachtigal falls |
| **101D2** | *Mormyrus* sp. "short snout" | 11.740 | -18.305 | dry | 19 | February | 2018 | Sanaga, Nachtigal falls |
| **101D3** | *Mormyrus* sp. "short snout" | 12.007 | -21.204 | dry | 19 | February | 2018 | Sanaga, Nachtigal falls |
| **101D4** | *Mormyrus* sp. "short snout" | 11.833 | -19.744 | dry | 19 | February | 2018 | Sanaga, Nachtigal falls |
| **101D6** | *Mormyrus* sp. "short snout" | 12.746 | -21.916 | dry | 19 | February | 2018 | Sanaga, Nachtigal falls |
| **101 E9** | *Mormyrus* sp. "short snout" | 12.504 | -19.260 | dry | 20 | February | 2018 | Sanaga, Nachtigal falls, upstream |
| **101F1** | *Mormyrus* sp. "short snout" | 12.265 | -20.308 | dry | 20 | February | 2018 | Sanaga, Nachtigal falls, upstream |
| **101F4** | *Mormyrus* sp. "short snout" | 12.266 | -21.895 | dry | 20 | February | 2018 | Sanaga, Nachtigal falls, upstream |
| **101F5** | *Mormyrus* sp. "short snout" | 12.187 | -22.424 | dry | 20 | February | 2018 | Sanaga, Nachtigal falls, upstream |
| **101F2** | *Mormyrus* sp. "short snout" | 11.988 | -18.628 | dry | 20 | February | 2018 | Sanaga, Nachtigal falls, upstream |
| **101F3** | *Mormyrus* sp. "short snout" | 11.806 | -19.274 | dry | 20 | February | 2018 | Sanaga, Nachtigal falls, upstream |
| **101F9** | *Mormyrus* sp. "short snout" | 12.547 | -21.291 | dry | 21 | February | 2018 | Avo'o |
| **101G1** | *Mormyrus* sp. "short snout" | 12.188 | -21.100 | dry | 21 | February | 2018 | Avo'o |
| **101G3** | *Mormyrus* sp. "short snout" | 12.054 | -20.109 | dry | 21 | February | 2018 | Avo'o |
| **101G4** | *Mormyrus* sp. "short snout" | 12.567 | -21.197 | dry | 21 | February | 2018 | Avo'o |
| **101G2** | *Mormyrus* sp. "short snout" | 12.295 | -23.053 | dry | 21 | February | 2018 | Avo'o |
| **13470** | *Paramormyrops batesii* | 11.351 | -26.533 | dry | 12 | February | 2018 | Stream park (Djim river) |
| **13679** | *Paramormyrops batesii* | 9.927 | -28.046 | dry | 13 | February | 2018 | Stream park (Djim river) |

| **code** | ***species*** | **d15N** | **d13C** | **Season** | **date** | **month** | **year** | **location** |
|---|---|---|---|---|---|---|---|---|
| **13023** | *Paramormyrops batesii* | 11.049 | -28.974 | dry | 6 | February | 2018 | Stream park (Mey river) |
| **12649** | *Paramormyrops batesii* | 10.338 | -27.146 | wet | 22 | November | 2017 | Stream park (Mey river) |
| **15881** | *Petrocephalus christyi* | 12.546 | -22.733 | wet | 14 | September | 2018 | Stream san (Nia river) |
| **62E1** | *Petrocephalus christyi* | 11.435 | -22.647 | dry | 9 | March | 2017 | Sanaga, Nachtigal falls |
| **62G6** | *Petrocephalus christyi* | 12.168 | -22.199 | dry | 11 | March | 2017 | Sanaga, Nachtigal falls |
| **62E4** | *Petrocephalus christyi* | 12.263 | -21.881 | dry | 9 | March | 2017 | Sanaga, Nachtigal falls |
| **62F7** | *Petrocephalus christyi* | 12.132 | -21.645 | dry | 10 | March | 2017 | Sanaga, Nachtigal falls |
| **14453** | *Petrocephalus christyi* | 10.978 | -32.835 | dry | 23 | March | 2018 | Stream san (Asamba river) |
| **14455** | *Petrocephalus christyi* | 11.530 | -27.179 | dry | 23 | March | 2018 | Stream san (Asamba river) |
| **14456** | *Petrocephalus christyi* | 11.577 | -27.098 | dry | 23 | March | 2018 | Stream san (Asamba river) |
| **14457** | *Petrocephalus christyi* | 10.257 | -26.428 | dry | 23 | March | 2018 | Stream san (Asamba river) |
| **15542** | *Petrocephalus christyi* | 11.877 | -28.995 | transition | 12 | July | 2018 | Stream san (Asamba river) |
| **13828** | *Petrocephalus christyi* | 11.321 | -33.540 | dry | 15 | February | 2018 | Stream park (Djim river) |
| **13829** | *Petrocephalus christyi* | 11.324 | -30.368 | dry | 15 | February | 2018 | Stream park (Djim river) |
| **13830** | *Petrocephalus christyi* | 10.295 | -36.892 | dry | 15 | February | 2018 | Stream park (Djim river) |
| **13831** | *Petrocephalus christyi* | 10.766 | -34.844 | dry | 15 | February | 2018 | Stream park (Djim river) |
| **13832** | *Petrocephalus christyi* | 11.352 | -35.719 | dry | 15 | February | 2018 | Stream park (Djim river) |
| **12827** | *Petrocephalus christyi* | 12.111 | -29.267 | dry | 5 | February | 2018 | Stream park (Mpem river) |
| **12828** | *Petrocephalus christyi* | 11.278 | -33.720 | dry | 5 | February | 2018 | Stream park (Mpem river) |
| **12829** | *Petrocephalus christyi* | 12.388 | -27.759 | dry | 5 | February | 2018 | Stream park (Mpem river) |
| **14192** | *Petrocephalus christyi* | 12.459 | -23.751 | dry | 20 | February | 2018 | Sanaga, Nachtigal falls, upstream |
| **14543** | *Petrocephalus christyi* | 11.223 | -29.351 | dry | 24 | March | 2018 | Stream san (Nia river) |
| **14544** | *Petrocephalus christyi* | 11.563 | -28.448 | dry | 24 | March | 2018 | Stream san (Nia river) |
| **14545** | *Petrocephalus christyi* | 11.299 | -26.535 | dry | 24 | March | 2018 | Stream san (Nia river) |
| **14679** | *Petrocephalus christyi* | 12.970 | -29.194 | dry | 27 | March | 2018 | Stream san (Sele river) |
| **15416** | *Petrocephalus christyi* | 13.152 | -27.259 | transition | 10 | June | 2018 | Stream san (Sele river) |
| **15417** | *Petrocephalus christyi* | 12.439 | -26.535 | transition | 10 | June | 2018 | Stream san (Sele river) |

| **code** | ***species*** | **d15N** | **d13C** | **Season** | **date** | **month** | **year** | **location** |
|---|---|---|---|---|---|---|---|---|
| **14767** | *Petrocephalus christyi* | 11.380 | -26.880 | dry | 28 | March | 2018 | Stream san (Tede river) |
| **14768** | *Petrocephalus christyi* | 12.288 | -32.244 | dry | 28 | March | 2018 | Stream san (Tede river) |
| **14769** | *Petrocephalus christyi* | 11.897 | -28.535 | dry | 28 | March | 2018 | Stream san (Tede river) |
| **14770** | *Petrocephalus christyi* | 11.899 | -24.655 | dry | 28 | March | 2018 | Stream san (Tede river) |
| **15482** | *Petrocephalus christyi* | 12.509 | -26.325 | transition | 11 | June | 2018 | Stream san (Tede river) |
| **15483** | *Petrocephalus christyi* | 12.435 | -26.295 | transition | 11 | June | 2018 | Stream san (Tede river) |
| **15485** | *Petrocephalus christyi* | 12.115 | -29.487 | transition | 11 | June | 2018 | Stream san (Tede river) |
| **15486** | *Petrocephalus christyi* | 11.962 | -33.244 | transition | 11 | June | 2018 | Stream san (Tede river) |
| **15487** | *Petrocephalus christyi* | 12.452 | -27.421 | transition | 11 | June | 2018 | Stream san (Tede river) |
| **101A1** | *Petrocephalus christyi* | 11.378 | -24.642 | dry | 18 | February | 2018 | Sanaga, Nachtigal falls |
| **101A2** | *Petrocephalus christyi* | 11.366 | -24.473 | dry | 18 | February | 2018 | Sanaga, Nachtigal falls |
| **101A3** | *Petrocephalus christyi* | 12.837 | -23.437 | dry | 18 | February | 2018 | Sanaga, Nachtigal falls |
| **101A5** | *Petrocephalus christyi* | 11.519 | -24.078 | dry | 18 | February | 2018 | Sanaga, Nachtigal falls |
| **101A4** | *Petrocephalus christyi* | 12.471 | -23.293 | dry | 18 | February | 2018 | Sanaga, Nachtigal falls |
| **101C5** | *Petrocephalus christyi* | 11.607 | -26.353 | dry | 19 | February | 2018 | Sanaga, Nachtigal falls |
| **101D1** | *Petrocephalus christyi* | 12.449 | -23.877 | dry | 19 | February | 2018 | Sanaga, Nachtigal falls |
| **101 E5** | *Petrocephalus christyi* | 12.126 | -23.541 | dry | 20 | February | 2018 | Sanaga, Nachtigal falls, upstream |
| **101 E6** | *Petrocephalus christyi* | 12.089 | -21.085 | dry | 20 | February | 2018 | Sanaga, Nachtigal falls, upstream |
| **101 E7** | *Petrocephalus christyi* | 12.476 | -21.758 | dry | 20 | February | 2018 | Sanaga, Nachtigal falls, upstream |

Table S2: sample list overview trophic samples

| sample | group | Amount | d15N | d13C | N% | C% |
|---|---|---|---|---|---|---|
| benthic | amphipoda | 0.49 | 7.26 | -24.98 | 10.00% | 37.50% |
| benthic | amphipoda | 0.512 | 7.10 | -24.96 | 6.89% | 31.57% |
| benthic | anisoptera | 0.536 | 6.99 | -27.05 | 11.54% | 46.08% |
| benthic | anisoptera | 0.542 | 6.49 | -22.04 | 11.11% | 43.57% |
| stomach content | anisoptera | 0.415 | 5.80 | -22.69 | 9.03% | 50.73% |
| stomach content | coleoptera | 0.522 | 7.20 | -30.58 | 10.77% | 48.42% |
| stomach content | coleoptera | 0.465 | 6.20 | -26.07 | 10.67% | 47.15% |
| benthic | coleoptera Hyrdophillidae | 0.532 | 5.81 | -21.12 | 11.74% | 47.83% |
| stomach content | coleoptera larvae Elmidae | 0.557 | 5.37 | -27.33 | 9.83% | 46.10% |
| stomach content | coleoptera larvae Elmidae | 0.506 | 5.47 | -24.90 | 9.80% | 46.29% |
| benthic | coleoptera larvae Elmidae | 0.488 | 3.94 | -19.96 | 9.66% | 46.44% |
| stomach content | coleoptera | 0.5 | 6.35 | -26.90 | 9.87% | 48.49% |
| stomach content | diptera | 0.337 | 6.84 | -26.24 | 10.95% | 45.08% |
| stomach content | diptera | 0.321 | 6.99 | -21.23 | 9.77% | 42.51% |
| stomach content | diptera | 0.535 | 7.32 | -16.68 | 11.39% | 44.38% |
| stomach content | ephemeroptera | 0.283 | 6.56 | -24.49 | 9.58% | 38.97% |
| stomach content | gastropoda | 0.52 | 5.94 | -17.82 | 1.99% | 7.12% |
| benthic | gastropoda | 0.489 | 6.77 | -20.03 | 2.92% | 21.55% |
| benthic | gastropoda | 0.47 | 6.56 | -18.06 | 1.54% | 16.16% |
| benthic | gastropoda | 0.516 | 6.23 | -14.77 | 1.05% | 14.77% |
| benthic | hemiptera | 0.536 | 7.92 | -25.19 | 11.70% | 47.75% |
| stomach content | lepidoptera | 0.461 | 6.15 | -24.04 | 8.49% | 42.58% |
| stomach content | lepidoptera | 0.471 | 6.70 | -23.89 | 9.34% | 45.27% |
| stomach content | lepidoptera | 0.452 | 7.31 | -21.60 | 9.59% | 48.76% |
| stomach content | lepidoptera pupa | 0.489 | 6.15 | -20.07 | 9.06% | 44.73% |
| benthic | odonata | 0.545 | 6.72 | -22.73 | 11.52% | 44.11% |
| benthic | odonata | 0.468 | 7.26 | -21.53 | 12.08% | 43.81% |
| benthic | plant | 2.528 | 7.55 | -22.99 | 4.02% | 36.91% |
| benthic | plant | 2.986 | 8.47 | -20.39 | 1.77% | 37.16% |
| stomach content | plecoptera | 0.31 | 4.83 | -25.74 | 9.12% | 43.90% |
| stomach content | trichoptera | 0.419 | 5.70 | -26.08 | 7.01% | 57.42% |
| benthic | zygoptera | 0.462 | 7.77 | -22.95 | 12.52% | 44.42% |
| benthic | zygoptera | 0.532 | 7.78 | -22.80 | 11.45% | 44.05% |

Table S3: results of single-factor ANOVAs and Tukey-Kramer test (if applicable) for species with transition season samples testing the factor of locality

| single-factor ANOVA | p-value | df | Tukey-Kramer q-stat values | dry vs wet | dry vs transition | wet vs transition | critical value with α=0.01 |
|---|---|---|---|---|---|---|---|
| *Mormyrus tapirus* δ13C without Mpem | 0.035* | 47 | | 2.591 | 2.400 | 3.415** | 2.678 |
| *Mormyrus tapirus* δ15N without Mpem | 0.109 | 47 | | | | | |
| *Marcusenius sanagaensis* δ13C without Mekono | 0.594 | 38 | | | | | |
| *Marcusenius sanagaensis* δ15N without Mekono | 0.765 | 38 | | | | | |
| *Marcusenius sanagaensis* δ13C without Mpem | 0.013* | 37 | | 0.455 | 3.840** | 4.223** | 2.719 |
| *Marcusenius sanagaensis* δ15N without Mpem | 0.013* | 37 | | 0.245 | 4.126** | 3.922** | 2.719 |
| *Petrocephalus christyi* δ13C dry vs transition without Mpem | 0.417 | 39 | | | | | |
| *Petrocephalus christyi* δ15N dry vs transition without Mpem | 0.011* | 39 | | | | | |

*p-value below 0.05 **q-stat value > critical value
